# JMod: Joint modeling of mass spectra for empowering multiplexed DIA proteomics

**DOI:** 10.1101/2025.05.22.655512

**Authors:** Kevin McDonnell, Daniel J. Geiszler, Nathan Wamsley, Jason Derks, Sarah Sipe, Zachary A. Cohen, Lina Kozhaya Warinner, Maddy Yeh, Eunice Koo, Andrew Leduc, Theodore J. Zwang, Harrison Specht, Nikolai Slavov

## Abstract

Parallelization of data acquisition substantially increases the throughput of mass spectrometry-based proteomics. However, parallelization also increases the density of mass spectra and consequently the overlap between ions, frustrating their analysis. To improve sequence identification and quantification from such spectra, we developed an open-source software for Joint Modeling of mass spectra (JMod). JMod models overlapping peaks as linear super-positions of their components in both MS1 and MS2 space, which permits multiplexed DIA with smaller mass offsets to increase the multiplexing capacity and thus proteomics throughput for a given plexDIA tag. This enables 9-plexDIA using 2 Da offset PSMtags, increasing throughput 9-fold while preserving quantitative accuracy and coverage depth. Furthermore, we use JMod to deconvolve simultaneous labeling by mass tags and heavy amino acids, thus increasing the throughput of metabolic pulse experiments measuring protein synthesis and degradation rates in single cells from mouse liver. By supporting enhanced decoding of highly multiplexed DIA spectra, JMod provides an open and flexible software that increases the throughput of sensitive proteomics.

## Introduction

Mass spectrometry (MS) is the most powerful and versatile method for protein analysis^1,2^. It enables deep and sensitive analysis down to single cells^3–8^ and single nuclei^9^, but its throughput remains limited^10,11^. Approaches that increase the number of peptides, samples, or both peptides and samples that can be analyzed in parallel can increase throughput but increase the density of mass peaks and thus pose computational challenges to the analysis of mass spectra.

In current acquisition schemes, data quality competes with sample throughput. Data-independent acquisition (DIA) achieves quantitative accuracy and the deepest proteome coverage without multidimensional chromatographic separation^12,13^. However, methods to increase throughput in DIA come with some constraints. Compressing separation times to achieve throughput exceeding 100 samples per day (SPD) significantly curtails peak sampling capacity and reduces quantification and coverage^10,12^. Since algorithms for analyzing DIA data^13–17^ have been a major contributor to expanding DIA performance, we reasoned that new computational workflows can further expand DIA capabilities, especially when tailored to new experimental designs and data acquisition approaches.

One such new approach is plexDIA. It achieves multiplexing in the mass domain using non-isobaric mass tags, where samples elute together, but their precursors and fragment ions can be uniquely attributed to their isotopically encoded labels^18–21^. However, natural peptide isotopic distributions will overlap if the channels’ mass differences are too small, effectively limiting tags to those which can provide at least 4 Da spacing between channels. Furthermore, experiments that depend on isotopic separation, such as metabolic pulse labeling^22^ or isoDTB activity-based protein profiling^23^, introduce additional overlapping isotopic envelopes that are incompatible with non-isobaric mass tags. These problems motivate the development of specialized tools to handle multiplexed data, especially the increased overlap between ions. This overlap increases both with the size of the isolation windows and with the number of multiplexed samples (Fig. 1**a**).

**Figure 1.**
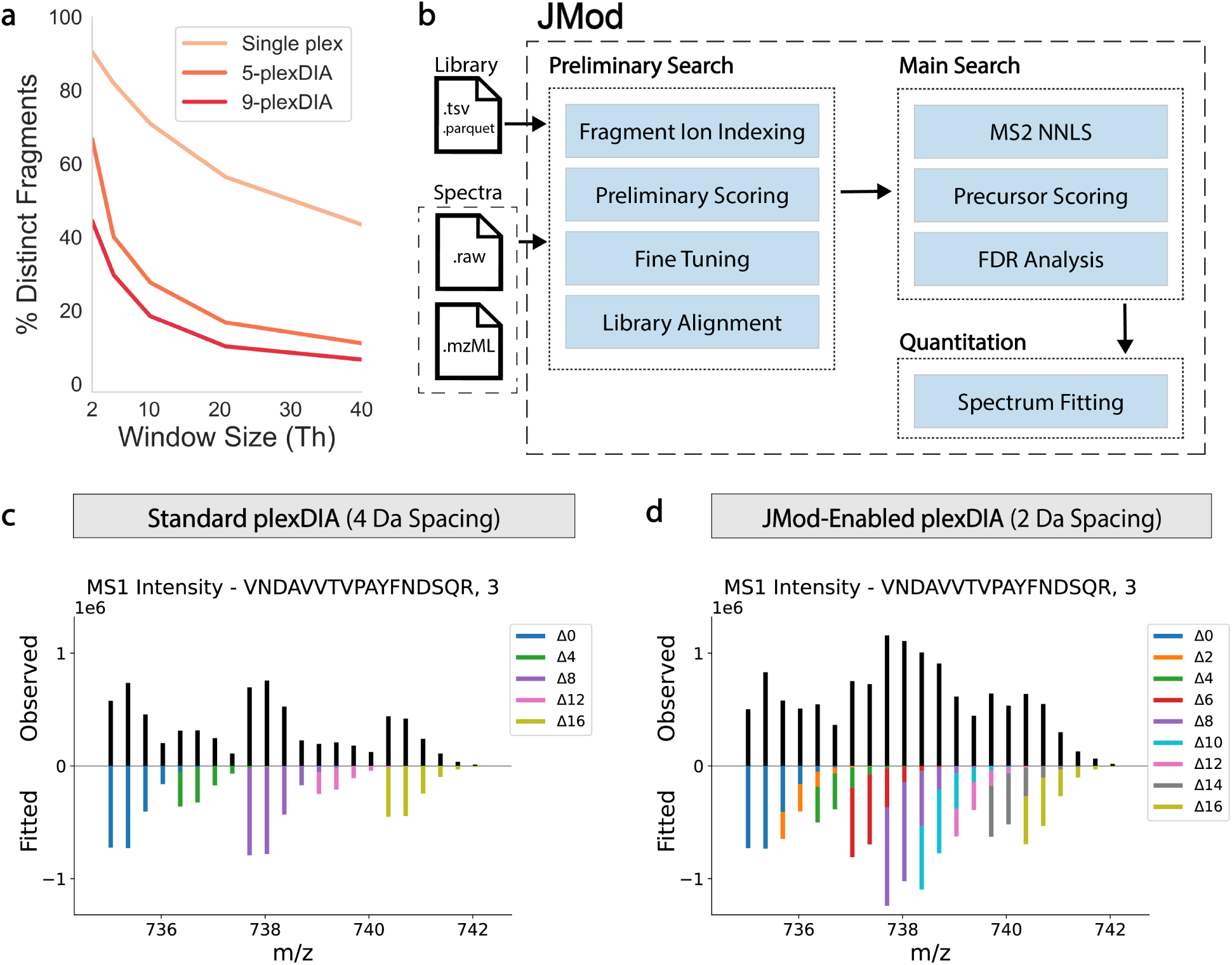
Modeling overlapping peaks in DIA mass spectra to increase multiplexing. **a** Median percentage of each precursor’s monoisotopic fragments that are distinct (not overlapping with other monoisotopic fragments) for single plex, 5-plexDIA, and 9-plexDIA across a range of window sizes. All fragments were derived from a library of 105,933 tryptic human precursors identified from 20 ng PSMtag samples as described in Methods. These precursors were then compared using a *m/z* tolerance of 0.01 Th and a retention time tolerance of 1.5% of the total range. y1 ions were not considered. **b** Schematic showing the major components of the JMod algorithm. **c** Mirror plot of the observed (top) and fitted (bottom) MS1 intensities for charge 3+ VNDAVVTVPAYFNDSQR precursor labeled with 5-plex PSMtags^29^ spaced by 4 Da. **d** Mirror plot of the same precursor but with 9-plex PSMtags spaced by 2 Da, uniquely enabled by JMod.

This overlap can undermine data interpretation unless addressed directly. A general and attractive approach for addressing it is to model the measured peak intensities as a linear superposition. Modeling mass spectra as a linear superposition was introduced decades ago^24^ and has been applied to model DIA fragmentation spectra by Specter^25^. However, it has not yet been implemented in robust tools that support DIA parallelization, such as plexDIA,^18,26,27^ and that achieve performance comparable to the current versions of DIA-NN and Spectronaut. Indeed, Specter did not support MS1 level quantification or the analysis of overlapping isotopic envelopes of precursors labeled with heavy isotopes.

To meet these challenges and enable increased multiplexing, we introduce JMod, an open-source software for DIA proteomics. JMod uses linear modeling to deconvolve overlapping peptide signals, allowing it to handle the increased complexity of multiplexed proteomics data while maintaining sensitivity and accuracy. Furthermore, JMod implements specialized routines to handle greater isotopic overlap in multiplexed DIA spectra than existing software, thus enabling greater sample parallelization for existing mass tags. In addition, as open-source software, JMod can serve as a community platform for the development of new computational workflows and capabilities in the field.

## Results

### The JMod algorithm

JMod is an open-source tool written in Python that can be run using a graphical user interface (GUI) or through the command line on any platform. It supports mass spectra provided in either .mzML or .raw format, and spectral libraries in .tsv or .parquet format. The algorithm can be broken down into three parts (Fig. 1**b**). First, a decoy-free preliminary search uses fragment ion indexing^28^ to identify a set of high-scoring target precursors with which to align the library to the observed data. The main search then fits the intensities of this updated library to the observed MS2 spectra using a non-negative least squares approach. The coefficients estimated by JMod from this fitting represent each precursor quantity for a given spectrum (Extended Data Fig. 1a). The data are then collapsed to candidate precursors and scored before being assigned a calibrated q-value. Finally, confidently identified precursors are quantified by performing additional fitting of the expected isotope distributions to the observed data. To support high quantitative accuracy, JMod introduces a new feature to the analysis of MS1 DIA spectra: Fitting the overlapping MS1 isotopic peaks across the elution profile, which results in a series of MS1 coefficients that define the MS1 elution peak of each channel. For more information, see Methods.

Since one element of JMod’s algorithm, the linear modeling of peptide fragments, has been used by Specter^25^, we sought to compare the performance of the two approaches. As Specter is no longer supported, we searched with JMod the data from the Specter article and compared the JMod results with the published Specter results. JMod identified over twice as many precursors from the same data at a lower false discovery rate (Extended Data Fig. 1b). This performance improvement is highly expected given the numerous differences and the algorithmic innovations in JMod (Fig. 1**b**).

### JMod enables increased multiplexing for the same mass tag

While analyzing mass spectra with overlapping peaks presents challenges, it can confer substantial benefits if the overlap is successfully modeled. Specifically, it enables an increase in the capacity for sample multiplexing as more distinct channels can be created for a mass tag using a set number of heavy isotopes. When plexDIA was introduced, it required 4 Da spacing between channels to minimize interference between channels^18^. The ability to relax this constraint can substantially increase sample throughput for many existing mass tags^30^.

As an intuitive visualization of this challenge and its solution, consider an example yeast peptide, multiplexed using PSMtag^29^ in the standard 4 Da spaced 5-plex configuration (Fig. 1**c**). The observed MS1 intensity is shown versus the fitted isotopes from JMod, with minimal overlap between the channels. The same peptide multiplexed with 2 Da spacing results in more complex, overlapping patterns (Fig. 1**d**). JMod deconstructs the spectrum into its constituent parts and thus uniquely enables plexDIA with the 2 Da spaced mass tags.

### Benchmarking sequence identification

To benchmark JMod, we first evaluated sequence identification on label-free samples containing mixed human and yeast proteomes using an entrapment library as it provides an objective measure of the false positive rate of identifications^31^. To replicate single-cell input amounts, human and *S. cerevisiae* tryptic protein digests were diluted to 200 pg and 80 pg respectively, per injection. These samples were analyzed on an Orbitrap Astral. We compared JMod with DIA-NN^13^, the state-of-the-art for processing both label-free and plexDIA data that is also freely available to the research community. In this experiment only, we used to DIA-NN version 1.9 as this was the latest version that returned an internal score (CScore) for every precursor which is required for entrapment analysis; all other comparisons use the latest version of DIA-NN, 2.5.1.

To evaluate both algorithms at the same stringency threshold, we plotted the number of human precursors and protein groups identified by each method as a function of the empirically derived entrapment false discovery rates (eFDR), (Fig. 2**a**). Although the inclusion of entrapment sequences in the library reduces the total number of identifications for both algorithms, it allows their relative performance to be fairly evaluated at a 1% eFDR^13^. The eFDR was calculated using the “paired method” as described by Wen *et al.*^31^, where every target sequence in the library has a corresponding shuffled entrapment sequence. In this analysis, JMod identified the same number of proteins (statistically indistinguishable) and 6% fewer precursors at 1% eFDR.

**Figure 2.**
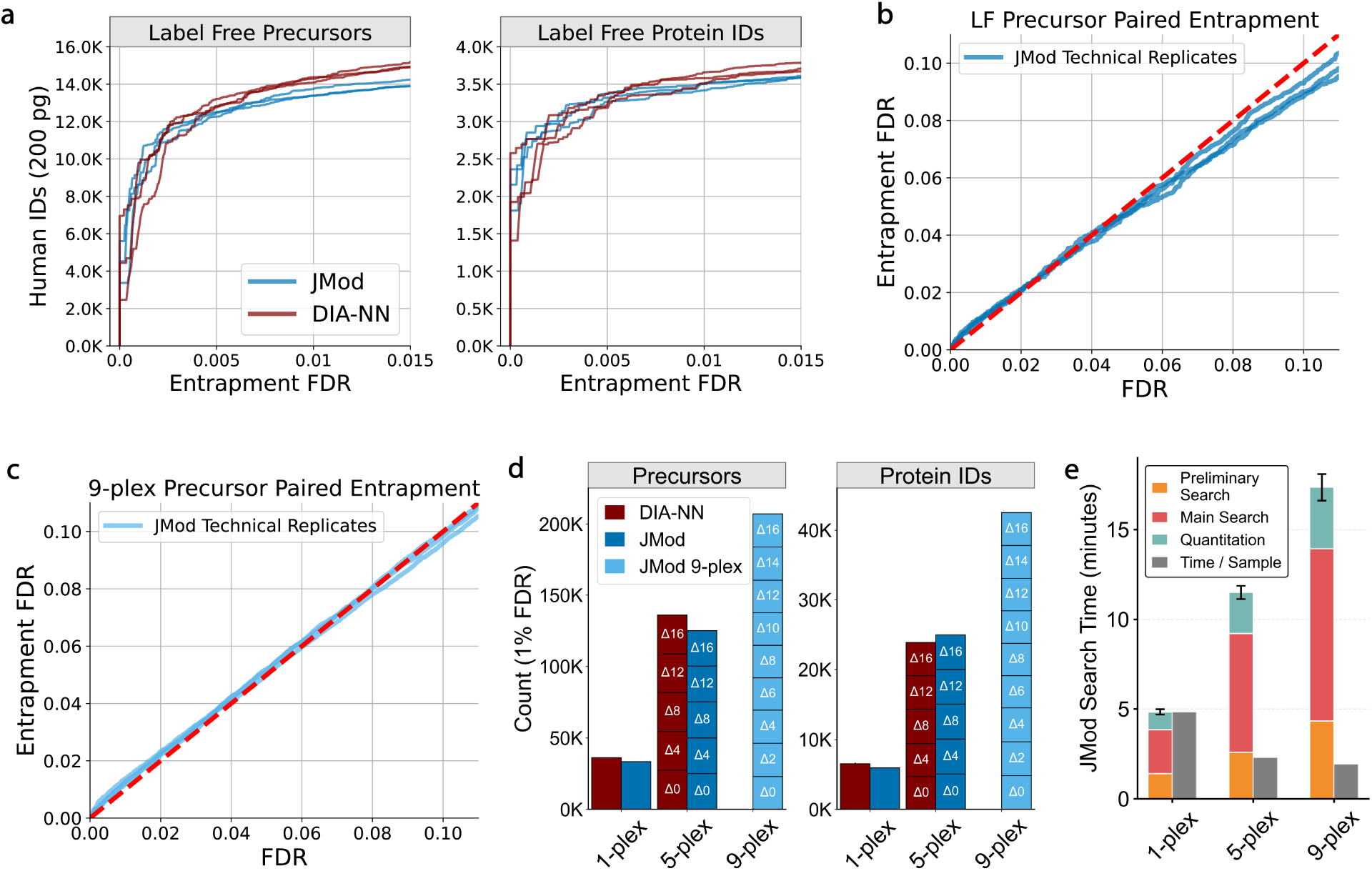
Evaluating sequence identification from label-free and multiplexed DIA data. **a** The identification performance of JMod (blue) and DIA-NN 1.9 (red) on a label-free sample composed of 200 pg human and 80 pg yeast peptides. The number of human precursors and proteins for each algorithm is plotted versus the entrapment FDR for 3 replicate injections. **b** The q-values reported by JMod versus the empirically derived q-values for precursors identified in the 200 pg label-free entrapment experiment. **c** The q-values reported by JMod versus the empirically derived q-values for precursors identified in the 200 pg per channel, PSMtag 9-plex entrapment experiment. **d** Human identification rates for JMod (blue) and DIA-NN 2.5.1 (red) from single and multiplexed PSMtag experiments. The 9-plex data were searched only with JMod because it uniquely supports plexDIA spectra with 2 Da channel offsets. **e** Bar plots showing the average search times for the JMod searches in the experiments from panel **d**.

We then sought to evaluate the validity of the q-values assigned by JMod to the identified precursors by comparing them to the eFDR (Fig. 2**b**). These q-values are typically estimated using internally generated decoys, which should reflect the distribution of false positive targets^32^. We compared the FDR estimates of JMod against the eFDR for 3 technical replicates of the label-free data. For these label-free data, JMod reported a well calibrated estimate of the FDR, shown by its proximity to the *x* = *y* (red dashed) line.

Next, we sought to evaluate the FDR control of JMod for highly multiplexed samples. Toward increased multiplexing, JMod enables, for the first time, searching DIA data from samples comprised of peptides with substantially overlapping isotopic envelopes. We therefore wished to show that the false discovery rate was controlled. For this evaluation, we used 9-plex replicates labeled with PSMtags^29^ which have channels separated by just 2 Da. The replicates contained mixed species samples described in Supplementary Table 1. Each sample contained 200 pg of human digest as well as *S. cerevisiae* digest ranging from 20 to 80 pg. We evaluated the search results using an entrapment analysis with shuffled sequences and the “paired method”. The results in Fig. 2**c** show that the empirically derived precursor FDR estimated by JMod is again an accurate representation of the eFDR for the 3 technical replicates of the 9-plex data, indicating that the reported false discovery rates are also well-calibrated for multiplexed data with overlapping isotopic envelopes.

### Proportionality of throughput to plex-size

As the number of samples analyzed in parallel using plexDIA increases, the number of quantified proteins per sample should ideally increase or remain the same, thus increasing the number of quantitative data points proportionally to the plex-size. We evaluated the ability of JMod to support such scaling with increasing plex-size using PSMtags^29^ across three different plex-sizes. Here, we also compared JMod to DIA-NN^13^ version 2.5.1 with QuantUMS^33^ (accuracy) as the quantification strategy. Single-cell input amounts of human and yeast proteomes were labeled with PSMtag and acquired in single-plex (Δ0 only), 5-plex (4 Da spaced) and 9-plex (2 Da spaced) schemes (Supplementary Table 1).

Both algorithms maintained the majority of their identifications as plex-size increased (Fig. 2**d**). DIA-NN identified 6,552 human proteins on average for the single-plex (Δ0), and 4,780 (73% of single-plex) for the 5-plex samples (Extended Data Fig. 2a). JMod identified 5,977 proteins on average from the single-plex data, and 4,999 (84% of single-plex) per channel for the 5-plex. Furthermore, JMod identified 4,724 (79% of single-plex) proteins on average from the 9-plex channels. This provided more than 40,000 human protein data points for each 9-plex run. These results indicate that JMod was better able to preserve the number of proteins quantified per sample as the plex-size increased. As previously noted, directly comparing identifications between search engines is difficult; q-value propagation across channels might be performed differently at both the precursor and protein level, and differences in how peptides are grouped into proteins can affect protein counts^34^. DIA-NN, for instance, reports protein q-values per channel rather than per run in plexDIA mode, which may be more conservative. For more information, see Methods.

### Decreasing search time per sample

With greater spectrum complexity, increased multiplexing can make the analysis of individual files harder. To show that the increased multiplexing enabled by JMod is actually computationally advantageous, we evaluated the run time of the algorithm across the PSMtag labeled data. These data were searched with an empirical library containing 55,453 precursors per channel and were collected at a rate of 30 runs per day (50 minutes per file) on an Orbitrap Astral with approximately 200 pg input per channel. Although the search time inevitably increased as the plex-size increased, the search time per sample decreased (Fig. 2**e**). JMod searched the 5-plex data at a rate 2.1 times faster per sample than the non-multiplexed data and the 9-plex data 2.5 times faster per sample. With the 9-plex searches taking less than 17.5 minutes, this equates to an analysis rate of greater than 740 samples per day. All of these data were analyzed using a MacBook Pro with a 14-core M4 Pro CPU and 48 GB of RAM.

### Accurate quantitation for overlapping isotopic envelopes

Next, we evaluated the accuracy of quantitation performed by JMod using the previously described mixing ratios of human and yeast proteomes. Proteomes mixed in different ratios at single-cell level input amounts were labeled with PSMtags (Supplementary Table 1) and analyzed by Orbitrap Astral. The ability of JMod to deconvolve the 2 Da spaced samples is exemplified by the yeast peptide TNNPETLVALR labeled with PSMtags in both the 5-plex and 9-plex schemes (Fig. 3**a**). Despite the additional overlap in the 9-plex configuration, JMod models the correct elution profiles for all channels as shown by the agreement of the MS1 coefficients for the 5 and 9 plex samples.

**Figure 3.**
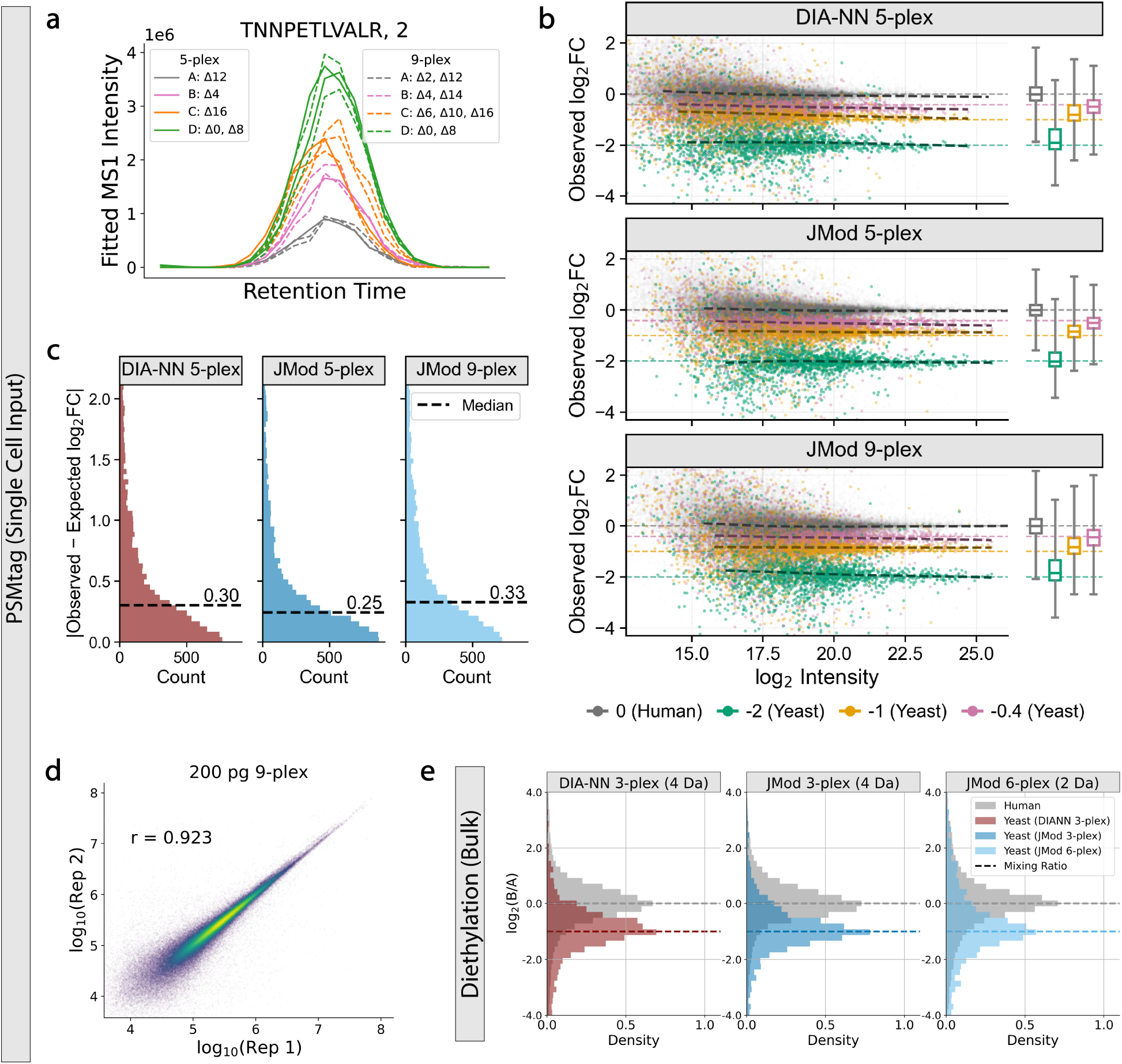
Evaluating quantitative accuracy for plexDIA using 4 Da and 2 Da offsets. **a** JMod’s fitted MS1 coefficients for the yeast precursor (TNNPETLVALR, 2+), in all channels from both a 5-plex and 9-plex MS run. Colors indicate the different mixtures and therefore input amount. **b** Scatter plots of the log_2_ fold changes (FC) versus the log_2_ intensity for intersected human and yeast precursors identified by JMod and DIA-NN 2.5.1 when searching the 5-plex PSMtag data as well as identified by JMod in the 9-plex data. Expected channel ratios are shown by the respective colored dashed lines while black dashed lines show the LOESS fit to each set of fold changes. Quantification by DIA-NN used the column *Ms1.Normalized* while quantification by JMod used the column *plex Area*. **c** Distribution of absolute errors between the observed ratios and the expected mixing ratios of intersected yeast precursors identified by JMod and DIA-NN 2.5.1 in the 5-plex PSMtag data and JMod in the 9-plex PSMtag data. **d** Scatter plot comparing JMod MS1 precursor quantities from two 9-plex replicates and their Pearson correlation. **e** Distributions of log_2_ fold changes for intersected diethyl labeled human and yeast precursors identified by both JMod and DIA-NN 2.5.1 when searching the 3-plex data separated by 4 Da, as well as identified by JMod when searching the 6-plex data separated by 2 Da. Data are from the Δ0 and Δ8 channels which have retention time separation. Expected ratios are shown by colored dashed lines. Quantification by DIA-NN used the column *Precursor.Normalized* while quantification by JMod used the column *plex Area*.

We then systematically compared the quantitative accuracy of both JMod and DIA-NN for these PSMtagged single-cell-level samples (Fig. 3**b**). To fairly assess the differences between the separate models and offsets, only precursors identified in all 3 analyses (DIA-NN 5-plex, JMod 5-plex, and JMod 9-plex) were used for the comparison. Furthermore, the algorithms were evaluated using MS1 level quantitation as this was the best performing method for both DIA-NN and JMod on these data. JMod was found to perform better than DIA-NN for the 5-plex data where the channels are separated by at least 4 Da. This is highlighted by the proximity of the median of the JMod’s estimated fold changes to the expected fold changes and the tightness of the interquartile range compared to DIA-NN. The accuracy of quantitation was very similar for JMod on the 9-plex data as indicated by similar medians, with slightly larger inter-quartile ranges for all fold changes. To better quantify the MS1 quantitative accuracy between the different algorithms and plex-sizes, we focused on the distributions of absolute fold change errors for yeast precursors shared between JMod and DIA-NN (Fig. 3**c**). JMod and the 4 Da spaced 5-plex data exhibited the highest accuracy, with the smallest median error of the three (0.25). The absolute accuracy of JMod on the 9-plex data separated by 2 Da (0.33), was similar to DIA-NN on the 5-plex data separated by 4-Da (0.30). A similar comparison for human precursors (Extended Data Fig. 2b) leads to the the same conclusion: JMod achieved the most accurate quantitation.

### Repeatability of highly multiplexed protein measurements

We then sought to evaluate the repeatability of these highly multiplexed measurements. Comparison between the precursors quantified by JMod from technical replicates of the 9-plex data show a Pearson correlation of 0.92 (Fig. 3**d**). To compare repeatability to DIA-NN, we next estimate the correlations between single-plex and 5-plex technical replicates analyzed both by DIA-NN and JMod (Extended Data Fig. 3). They achieved similar respective correlations: 0.98 and 0.96 for DIA-NN and 0.98 and 0.95 for JMod. We also evaluated the coefficients of variation (CV) for both precursors and proteins identified by JMod in highly multiplexed data by comparing them to those of label-free data. We compared the CVs of the identifications from the three replicates of the 200 pg per channel 9-plex data to those identified in three replicates of 200 pg label-free runs by DIA-NN and JMod (Extended Data Fig. 4). For the 2,568 shared protein identifications, JMod was able to provide similar, albeit lower, repeatability for the 9-plex channels compared to the label-free data. Proteins identified in the 9-plex data had median CVs between 0.07 and 0.1, compared to a median of 0.04 for both DIA-NN and JMod on the label-free data. Together, these results indicate that JMod can produce repeatable and precise quantitative performance for overlapping plexDIA channels.

Furthermore, previous studies have shown the advantages that plexDIA provides in terms of data completeness, but only for smaller plex-sizes^18^. To evaluate this in our experiments, we compared the reproducibility of the identifications in the highly multiplexed PSMtag data for both algorithms (Extended Data Fig. 5). JMod exhibited high data completeness in both the 5-plex and 9-plex data, Jaccard similarity greater than 0.97 across channels and greater than 0.84 across runs for both experiments. DIA-NN performed similarly with respective values of 0.95 and 0.84. This suggests that JMod can also provide consistent and repeatable identifications as well as quantitation.

### Generalizability of JMod across instruments and mass tags

To demonstrate the ability of JMod to generalize across mass tags and MS instruments, we evaluated the quantitative accuracy of JMod when searching diethylated bulk samples separated by both 4 Da and 2 Da. Diethlyation can provide multiplexing capabilities of 6-plex if allowing just 2 Da offsets between channels. However, deuteration of the diethyl tags introduces retention time shifts between mass channels that complicates quantitation^35^.

We again performed a mixed species experiment with mixed human and yeast proteomes, this time using an Exploris 480 instrument. Bulk mixed human and yeast proteomes were labeled using diethylation with both a 4 Da spaced 3-plex scheme and a 2 Da spaced 6-plex scheme (Supplementary Table 2). The evaluation was performed using the best quantitation method for both algorithms, with *Precursor.Normalised* and *plex Area* being used for DIA-NN and JMod respectively. Comparing the measured precursor ratios to the mixing ones shows JMod again achieving higher quantitative accuracy to DIA-NN 2.5.1 on 4-Da-spaced data, the now 3-plex, diethylated samples (Fig. 3**e**). Furthermore, JMod alone can quantify the 6-plex diethylation scheme relying on 2-Da spacing. Since JMod does not require channels to elute at the same time, it can model the isotopic overlap across the elution of all channels despite the deuterium retention time shifts (Extended Data Fig. 6a). The partial overlap can therefore be modeled with the area under the MS1 coefficients still providing accurate quantitation.

### Quantifying large fold changes with plexDIA

The accuracy of quantitation is typically evaluated for changes that are smaller than 5-fold, while biological samples may exhibit a much wider range of variation. Such wider range of variation can be particularly challenging to quantify when multiplexing with less than 4 Da and modeling overlapping mass peaks. To investigate how far we can achieve accurate quantitation in such scenarios, we performed additional quantitative evaluations of highly multiplexed data, this time with much larger fold changes between channels and across various on-column loads. Specifically, we evaluated ratio compression and identification in mixed-species benchmarks with spike-in abundances spanning 1- to 32-fold changes (Fig. 4). This dynamic range was evaluated for input amounts of 20ng (**a**-**c**), 2ng (**d**-**f**), and 200pg (**g**-**i**) per channel. The samples consisted of constant amounts of human protein digest and variable amounts of yeast (Supplementary Table 3) that were both labeled with PSMtag and acquired on an Orbitrap Astral mass spectrometer. We then compared the performance of JMod to DIA-NN 2.5.1 again with QuantUMS^33^ enabled and set to high accuracy to minimize ratio compression. To avoid any biases arising from differences in identifications, for a precursor to be considered in any quantitative comparisons, it must be quantified in the same channel in DIA-NN for MS1 and QuantUMS, JMod 5-plex, and Jmod 9-plex datasets.

**Figure 4.**
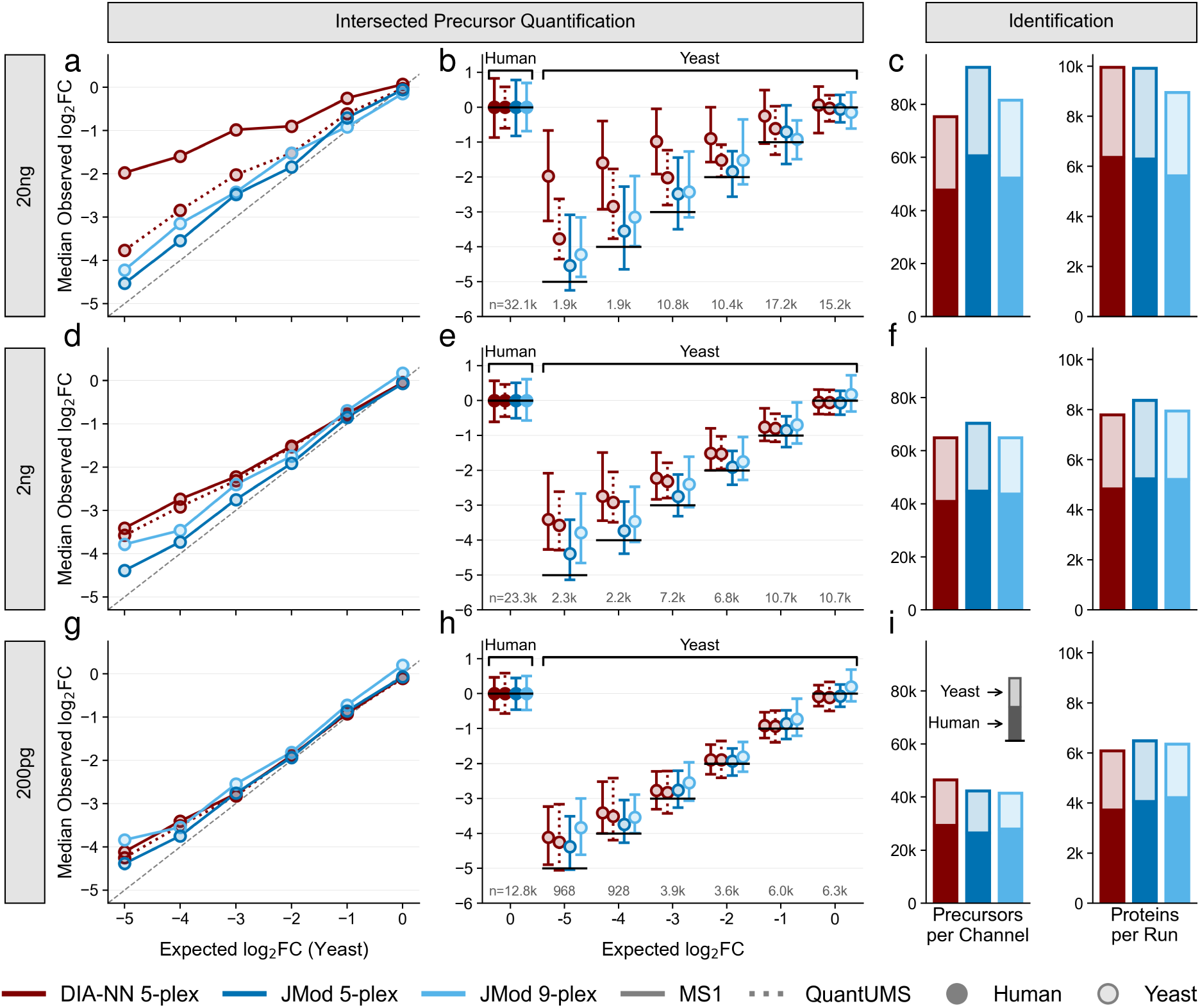
Evaluating large fold changes across a range of input amounts. a, d, **g** Median observed log_2_ fold changes (FC) versus the expected log_2_ FC for intersected precursors between JMod and DIA-NN 2.5.1 across per-channel input amounts of 20 ng, 2 ng, and 0.2 ng respectively. **b, e, h** Inter-quartile range about the median of log_2_ FCs versus the expected log_2_ FC for intersected precursors between JMod and DIA-NN 2.5.1 across per-channel input amounts of 20 ng, 2 ng, and 0.2 ng. **c, f, i** Total number of precursors and proteins identified per channel across input amounts of 20 ng, 2 ng, and 0.2 ng, for DIA-NN 2.5.1 and JMod on 5-plex data and JMod only for 9-plex data collected. In all panels, for MS1 quantification by DIA-NN we used the column *Ms1.Normalized* and for QuantUMS we used the column *Precursor.Normalized* from its output. For JMod all quantification shown is at MS1 levels using the column *plex Area* from its output.

At 20ng of human sample per channel, JMod exhibits less ratio compression than DIA-NN regardless of whether DIA-NN’s MS1 or QuantUMS quantification is used (Fig. 4**a**). Notably, this finding extends even to the 9-plex data, for which JMod quantifies about 734k precursors per experiment. This indicates that increasing sample throughput is consistent with high quantification accuracy even at bulk levels. We also observe no meaningful differences in data spread between methods at the largest fold changes (Fig. 4**b**), although at smaller fold changes JMod achieves tighter estimates than DIA-NN’s MS1 quantification and wider estimates than DIA-NN’s QuantUMS quantification. This is expected, as the increased specificity of fragment ions at the MS2 level provide cleaner signal–assuming the fragment ions are actually present–than the more crowded MS1 spectra. Finally, JMod is shown to achieve rough parity with DIA-NN on precursor and protein identifications (Fig. 4**c**). As mentioned previously, direct comparisons between protein identifications are difficult due to differences in how run-level q-values are calculated, so these values are likely to be a conservative estimate of DIA-NN’s protein identifications.

At lower input amounts, the ratio compression differences between DIA-NN’s MS1 and QuantUMS quantification diminishes, yet JMod is consistently found to be more accurate than DIA- NN. For example, at 2ng input, JMod estimates the 32-fold change (log_2_ fold change = 5) to be roughly 23-fold, while DIA-NN’s best estimate is closer to 11-fold (Fig. 4**d**); this corresponds to quantification of the lowest abundance precursors in the sample being 2-fold more accurate when quantified using JMod with the same level of measurement consistency (Fig. 4**e**). Although DIA-NN’s accuracy does begin to converge with JMod at the lowest input amounts when performing QuantUMS quantification (200pg, Fig. 4**g**), JMod is able to recover more consistent quantification values across the spectrum of fold changes (Fig. 4**h**). While JMod quantifies all precursors at MS2 levels, the benchmarks in Fig. 4 rely only on the MS1 level quantification, which supports accurate ratio estimation even when using 9-plex tags all the way up to 1 : 2^5^ fold changes, the largest dynamic range measured. This is particularly notable because the experiment design had the largest and smallest fold changes directly adjacent, indicating that even in the case of systematic 32-fold differences in intensity with overlapping isotopic envelopes JMod is able to recover reasonably accurate values.

Overall, this analysis reveals several things. First, JMod’s modeling of the isotope overlap can reduce MS1-based ratio compression even at high input amounts. Second, JMod achieves more accurate estimates of precursor quantity in our benchmarks. Finally, JMod is able to maintain accuracy and consistency of quantification with 2 Da-spaced reagents even at quantitative extremes.

### Increasing the throughput of metabolic-pulse single-cell proteomics

Metabolic pulse labeling of single cells has emerged as powerful technique to study the regulation of protein degradation^6,22^. However, because it requires utilizing isotopically doped amino acid feed, the associated isotopic envelopes would overlap and confound quantification of plexDIA-style mass tags. In practice, when combining heavy +6 Da lysine with plexDIA, channels would typically require at least 10 Da spacing to avoid isotopic overlap^22^. When multiplexing with mTRAQ, this limits the number of samples to 2 per run. Thus, without being able to model the overlapping isotopic envelopes, the throughput remains limited. By modeling the isotopic overlap, JMod should be able to enable 3-plex mTRAQ with metabolic labeling, thus increasing sample throughput by 50% over previous experiments.

Two test and exploit this possibility, we fed two mice +6 Da heavy lysine for 3 or 5 days. Then, their hepatocytes were extracted and labeled with mTRAQ (Fig. 5**a**). We used either 2-plex mTRAQ (as a control) or 3-plex. With this experimental design, JMod is able to model and quantify distinctly the light and heavy peptides even for the 3-plex (Fig. 5**b**). To evaluate the accuracy of this analysis, we first compared measured values between the 2-plex and 3-plex configurations. JMod identified similar numbers of proteins per cell in both plexes, with 1,016 proteins on average from 2-plex hepatocytes and 1,143 proteins on average from 3-plex hepatocytes (Fig. 5**c**).

**Figure 5.**
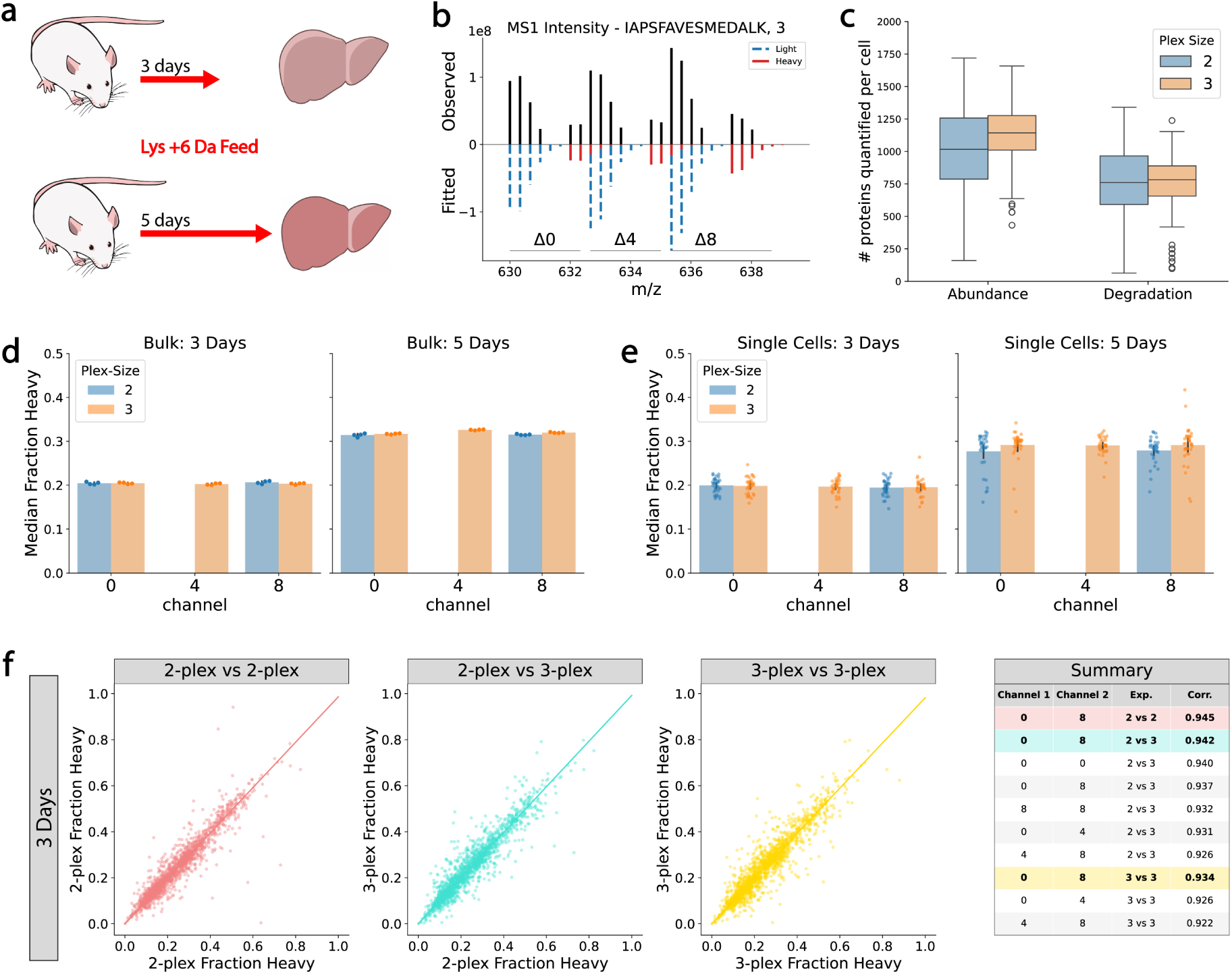
Comparing 4 Da and 2 Da spaced plexDIA combined with metabolic labeling. **a** Two mice were fed with +6 Da heavy lysine for 3 or 5 days. The livers of the mice were then harvested for analysis. **b** Example mirror plot of the observed and fitted MS1 intensities for the mTRAQ and metabolically labeled precursor IAPSFAVESMEDALK 3+. Lysine containing peptides contain both heavy and light versions, which can overlap with the next mTRAQ channel. JMod is uniquely able to deconvolve the signal as shown. **c** Box plots showing the numbers of proteins identified in single cells that were analyzed in either a 2-plex (4 Da spaced, blue) or 3-plex (2 Da spaced, orange) configuration. Total protein numbers (Abundance) as well as proteins with lysine-containing peptides (Degradation) are both shown. **d** Median fraction heavy of all precursors in bulk replicates acquired in both 2-plex and 3-plex schemes. **e** Median fraction heavy of all precursors in single hepatocytes acquired in both 2-plex and 3-plex schemes. **f** Example comparisons of the fraction Heavy for all precursors averaged per channel across equal numbers of single cells between the 2-plex and 3-plex acquisition schemes. Summary table shows the Pearson correlations between all possible comparisons.

We then evaluated the ability of JMod to maintain accurate quantitation between 2-plex and 3-plex configurations both for bulk and for single-cell measurements. This was benchmarked by the fraction of heavy peptides: heavy / (heavy + light). The median of this fraction is statistically identical for the 2-plex and 3-plex bulk samples (Fig. 5**d**). The heavy fraction was used for the comparison as this ratio includes measurements from different overlapping channels meaning it should highlight any quantification issues if they are present. In both mice (3 and 5 days), we see high agreement between the replicates across conditions with no observable difference between the data with overlapping isotopic envelopes (3-plex) and without (2-plex). Single cells were then evaluated in the same way (Fig. 5**e**). Again, JMod provides high agreement between the median fraction of the heavy channel for all precursors in cells acquired with the 2-plex and 3-plex configurations. Moreover, the results also agree with the bulk data, with a larger spread of values due to the single-cell variability.

Finally, we investigated the measured data for biases due to overlapping isotopic envelopes by looking at the average heavy fraction for each individual precursor across the single cells. Comparisons of the average heavy fraction per precursor across all possible plex combinations indicate excellent agreement and no evidence of bias (Fig. 5**f**). The corresponding Pearson correlations are greater than 0.92 for all comparisons, as shown in the summary table of Fig. 5**f**.

### Protein measurements define liver’s central portal axis

Projecting all collected hepatocytes onto their principal components (PC) reveals that the largest source of variance (PC1) is strongly correlated with the abundance of the liver zonation marker *Gulo*^36^ (Fig. 6**a**). To test if PC1 corresponds to the zonation axis, we examined how the 10 most differential portal and central markers of liver zonation^36^ vary in our single cells based on PC1 (Fig. 6**b**). Central markers show a strong positive correlation to PC1 indicating that the axis runs portal to central as indicated by Fig. 6**a**. Portal markers are strongly negatively correlated to PC1, again validating this result. Interestingly, the portal markers show a “hockey-stick” curve which has previously been established in spatial proteomics experiments^37^. This indicates that the accuracy of quantitation and therefore the biological validity provided by JMod from dissociated tissue is such that it can provide spatial resolution similar to that of cells acquired through laser microdissection.

**Figure 6.**
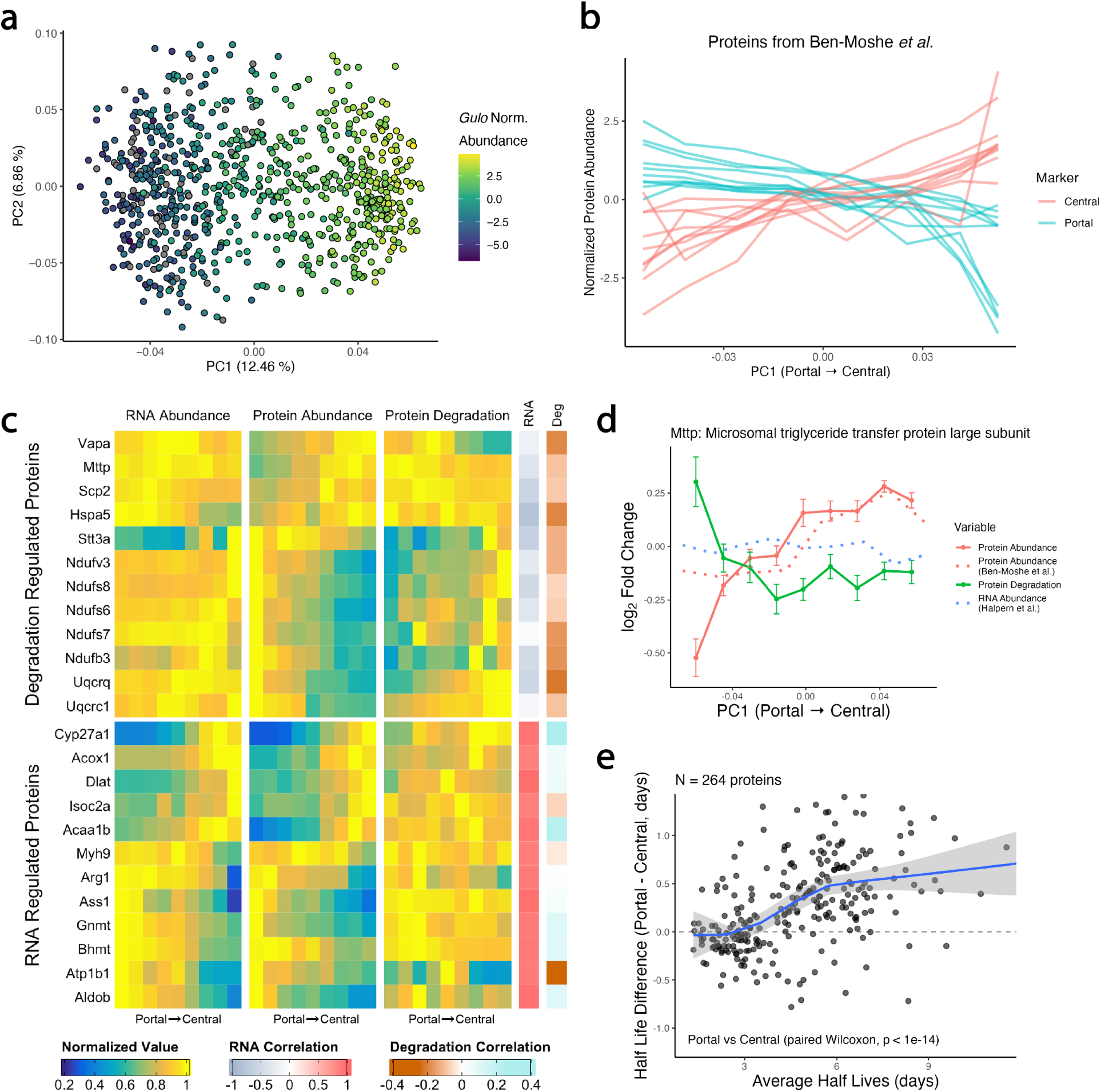
Quantifying protein abundance, synthesis and degradation in single hepatocytes reveals regulatory trends across the central portal axis. **a** PCA of the multiplexed hepatocytes showing the measured protein abundance of the central marker *Gulo*. **b** Single cells were sorted based on PC1 and the averaged abundance of protein markers^36^ displayed. **c** Comparison of protein abundance, protein degradation, and RNA abundance^38^ for hepatocytes across the axis of zonation. Proteins were subset to the 10 with the strongest negative correlation between degradation and abundance and the 10 with the strongest positive correlation between RNA and protein abundance. All proteins were required to be present in at least 300 cells. Correlations across the axis of zonation are shown between RNA abundance and proteins abundance (RNA Correlation), as well between protein degradation and protein abundance (Degradation Correlation). **d** Example of a protein (*Mttp*) whose abundance across the portal-central axis is inversely correlated to the rate of degradation but not to RNA abundance. Also shown are protein abundance and RNA abundance from previous studies^36,38^. **e** Comparison of the differences in half-lives of 264 proteins as measured in portal cells versus central cells, plotted against the average measured half life of the protein. Cells were labeled portal or central based on which tertile of PC1 they were in (lower tertile for portal, upper tertile for central). Only proteins that were present in at least half of all portal cells and half of all central cells were considered.

### Protein regulation across liver’s central portal axis

Having confirmed that we can accurately infer the zonation of the extracted hepatocytes, we looked to see if we could identify zonation proteins whose abundance was regulated by degradation. Previous studies have examined the correlation of RNA and protein abundance across the zonation of hepatocytes^36,38^, but we are not aware of studies examining how protein degradation impacts zonation at the single-cell level.

To do so we looked at RNA abundance, protein abundance, and protein degradation rates relative to the axis of zonation for proteins where abundance and degradation were strongly negatively correlated (Fig. 6**c**). RNA abundance was taken from the fluorescence-activated cell sorting (FACS) cells from Halpern *et al.*^38^, while protein abundance and degradation rates are from the JMod analyzed hepatocytes. For many proteins shown in Fig. 6**c**, RNA does not correlate with the protein abundance, while degradation does. This suggests that their zonation may be driven by the degradation rate. To complement this we also examined the proteins where RNA strongly correlates with the measured protein abundance. For these proteins, degradation was not found to negatively correlate as these proteins are regulated by RNA.

To further examine the role of protein regulation, we plotted the protein abundance, protein degradation rate and RNA abundance of *Mttp* (Fig. 6**d**). The protein abundance increases from portal to central, in line with a previous measurement by Ben-Moshe *et al.*^36^. This trend may be explained by a parallel decrease in its degradation rate, which can explain its acculturation towards the central zone. This role of degradation is bolstered by the measured RNA abundance of *Mttp* from Halpern *et al.*, which shows no correlation with the abundance of the protein. Under steady state the synthesis rate can be calculated as the protein abundance times the degradation rate^22^. Estimating this for *Mttp* shows an unchanging rate across the portal-central axis, again suggesting the zonation is driven by the rate of degradation.

This contribution of protein degradation (*K_deg_*) to establishing spatial protein gradients is confirmed by systematic analysis of half lives (*log*(2)*/K_deg_*) across all GO-terms (Extended Data Fig. 7). The results show many significantly zonated proteins. Importantly, GO-terms that showed significant zonation of half lives exhibited corresponding significant zonation of protein abundance.

Interestingly, we observed a systematic increase in the stability of proteins in portal cells. Specifically, the longer the life time of a protein is, the more its stability increases from central to portal cells (Fig. 6**e**). Portal cells exhibited longer half-lives for proteins present in at least half of all cells (N = 264, *p <* 10*^−^*^14^). As portal cells are closer to arteries, we verified that this was not a technical artifact due to the availability of the heavy lysine. Both portal and central cells showed similar availability of heavy lysine in their free pool (Extended Data Fig. 8).

## Discussion

Biomedical research needs scalable and affordable analysis of many thousands of samples across diverse applications ranging from population-scale plasma to single-cell proteomics^39–41^. A promising avenue to achieve such scalability is to parallelize data acquisition^11^. However, such approaches increase data density and lead to increased frequency of overlapping MS1 and MS2 signals (Fig. 1**a**). To model these challenges, we developed JMod, an open-source platform that can support scaling of MS proteomics by parallelization.

JMod seeks to support all forms of parallelization by modeling the predictable structure of multiplexed DIA data. Here, we demonstrate unique support for increased overlap between channels of non-isobaric mass tags. This allows the use of smaller mass offsets and therefore enables about 2-fold higher throughput with PSMtags and diethylation. Furthermore, it supports increased throughput of multiplexing in vivo metabolic pulse experiments multiplexed with mTRAQ. Similarly, JMod supports sample parallelization in time by introducing small delays between sample injections on the same instrument that produce overlapping active gradients. This method, termed timePlex^27^, inherently produces more crowded spectra that are amenable to the joint modeling technique and is directly and uniquely enabled by JMod.

Beyond supporting highly parallelized data acquisition, JMod offers advantages for the analysis of peptides with similar amino acid sequences as it can model their overlapping fragments as shown in Extended Data Fig. 1a. This advantage should benefit co-eluting peptides from post-translationally modified proteins and proteoforms originating from alternate RNA decoding^42^. Such modeling of overlapping peptide fragmentation spectra was first prominently used for DIA by Specter^25^ and more recently by Chimerys^16^ and Pioneer^43^. However none of these packages can support increasing throughput by plexDIA tags with small mass offsets, which is a primary objective of JMod. Further, JMod’s approach to quantification achieves high accuracy over a wide range of protein changes (Fig. 4) to the point of exceeding the quantitative performance of DIA-NN.

JMod can also support increasing levels of multiplexing alongside metabolically incorporated heavy amino acids offset by smaller mass differences. This can increase the throughput of quantifying protein synthesis and degradation rates^44,45^, thereby identifying mechanisms for liver zonation (Fig. 6**c, d**). Our results demonstrate a strong correspondence between protein abundance and protein degradation, thus showing how variation in protein degradation rates contributes to establishing spatial protein gradients (Fig. 6 and Extended Data Fig. 7).

The joint modeling approach of JMod provides highly accurate quantitation of precursors from single-cell-level input amounts to bulk, regardless of the channel overlap (Fig. 4). The quantification of 734k precursors in a single 9-plex set of 20 ng samples marks a promising data point for the ability of MS to scale throughput by multiplexing to accurately quantify many precursors per run^11,19,30^. While we benchmark accurate quantification using only MS1 level orbitrap data, JMod quantifies all precursors at MS2 level as well. Future versions can combine both MS1 and MS2 level data to further improve quantification. For 4 Da spaced channels, which are supported by other software, JMod performed better than the state-of-the-art software DIA-NN 2.5.1. This trend was consistent over the range of fold-changes and input amounts tested, highlighting the robustness of JMod’s quantitation and the feasibility of deep and accurate DIA quantification using only MS1 scans. This greater accuracy will not only serve to increase our ability to find biological differences between samples but is also essential to quantitative modeling and causal inference^26,46^. The use of tags with 2 Da offsets offers advantages and limitations. For the given molecular composition of the tags, JMod’s support for reduced mass offsets increases the maximal plex that can be designed. Furthermore, the overlapping isotopes decrease the total number of peaks per spectrum for a given plex, which reduces the potential for interference by close peaks compared to the same plex achieved with 4 Da mass tags. While 2 Da plexDIA spacing does enable increased throughput, the increased measurement errors can slightly reduce the ability to quantify very large fold changes (Fig. 4**a**). However, quantitative accuracy was still high across all JMod evaluations for both spacings, with JMod accuracy with 2 Da spacing often outperforming other methods with 4 Da spacing.

We expect to continue building new features into JMod for enhancing sequence identification and quantification from highly parallelized DIA data. These features will support new experimental designs, as in the case of timePlex, support more data types (including ion mobility), and further exploit the inherent and engineered structure of multiplexed data^19^. We also hope that the open-source nature of JMod will encourage others to contribute to these future developments.

## Data and Code availability

The source code for JMod is available at: github.com/ParallelSquared/JMod under the Apache License 2.0. Raw data, spectral libraries, search outputs, and meta data are available at MassIVE: MSV000097957 and via ProteomeXchange with identifier PXD083370. The code used to generate the figures is available at: github.com/ParallelSquared/JModFigures

## Acknowledgments

We thank O. Kok for help with single-cell sample preparation; V. Demichev for providing feedback on running DIA-NN optimally; R. Thomas for maintaining the Astral at top performance; and the members of Parallel Squared Technology Institute for their constructive feedback.

## Author Contributions

**Software and algorithmic design**: K.M.D, D.J.G., N.W., J.D., Z.C., and E.K. **Experimental design**: K.M.D., H.S. and N.S **Sample preparation and acquisition**: S.S., L.K.W., M.Y., and T.J.Z. **Data analysis**: K.M.D., D.J.G., N.W., J.D., and A.L. **Writing & editing**: K.M.D., D.J.G., N.W., S.S. and N.S. **Raising funding & supervision**: H.S. and N.S.

## Methods

### The JMod Algorithm

#### Data input and processing

JMod currently accepts centroided data in .mzML and .raw format and libraries in tab delimited or .parquet format. If specified, JMod will add additional fragment isotopes to the library using the BrainPy package^47^.

#### Preliminary search

JMod uses a preliminary search to calibrate spectra, align library and experimental retention times, and optimize m/z and retention time tolerances. This helps account for differences between library and experimental gradients as well as errors in measured peak *m/z*. The preliminary search is performed using the internally developed peppy sage Python library that is forked from Sage^48^. This implements a fragmention indexing approach that restricts the search space and ranks candidates by hyperscore based only on fragments that are present in the provided spectral library. MS2 spectra closest to the apex of each feature are searched for possible library candidates matching the precursor *m/z* using a wide (20 ppm) tolerance. These identifications are then sorted by how well they fit based on their Scribe score^49^. JMod iteratively restricts to higher quality identifications, stopping when the retention time alignment errors can be fit with a Gaussian mixture model where 99% of the mass is attributable to the true positive distribution. This step removes low-scoring false positives, preventing them from impacting the alignment.

#### Retention time alignment and fine-tuning

JMod also employs machine learning to fine-tune the library to the experimental conditions using the precursors identified in the preliminary search. This is similar to an approach introduced by Wallmann *et al.*^50^. JMod then uses the peptide sequences of high-scoring precursors and their observed retention times to fine-tune its internal, pretrained, convolutional neural network (CNN) models. These models take an input of one-hot encoded amino acids and predict the scaler retention time. If enough peptides are found, the models are fine-tuned using the training data for 50 epochs or fewer as this step includes early-stopping to prevent overfitting. Fine-tuning provides much better predictions of observed retention times, allowing JMod to use a much stricter retention time tolerance. Depending on the experiment, JMod automatically selects one of four different base models: label free, mTRAQ, diethylation, and PSMtag. A user can also fine-tune a library on their own data and use this for their subsequent searches instead of the base models. This is particularly useful for low-input samples, where the preliminary search may not recover enough IDs to perform adequate fine-tuning.

Once fine-tuning is complete, both the updated predictions for the validation set and the original library retention times are compared to observed values. JMod will select the fine-tuned predictions; only if they better describe the data. Otherwise, the original retention times will be used. More specifically, the area under the cumulative distribution of the absolute retention time errors is used to decide on the best model. Those with smaller errors will have a larger area under this curve. The error distributions are modeled as both Gaussian and exponential distributions, with the best approximation being used to derive optimized retention time tolerances. Users can also skip the fine-tuning step if they know that the library contains accurate empirical retention times.

The high-scoring precursors are also used to correct for any systematic *m/z* errors. A LOWESS regression is first used to correct for changes in *m/z* error across retention time. A subsequent LOWESS regression line is used to account for *m/z* errors that depend on the *m/z* itself. An *m/z* tolerance is then calculated as 4 standard deviations of the Gaussian distribution fit to the MS1 errors from the preliminary search.

#### Main search

Before the main search, the library retention times and precursor *m/z* are adjusted to match the experiment using the models derived in the preliminary search. If performing plexDIA or timePlex experiments, the library is also duplicated for each channel, with each receiving unique retention times and/or *m/z* values; JMod natively supports multiple labeling schemes including PSMtag, mTRAQ, diethylation, and metabolic labeling, while users can also add custom tags to the tag library. Decoys are generated for each precursor in the library using a pseudo-reverse method^32^, with the data then searched using the combined target and decoy libraries.

During the main search, JMod models each MS2 spectrum independently as a linear super-position of the library precursors. Library precursors that fall within the retention time tolerance and window *m/z* range are considered as potential candidates. Those candidates that do not match greater than 20% of their fragment intensity are then excluded. These parameters can be adapted to the experiment in question if required. The remaining precursors are used to set up non-negative least squares problem in the form:

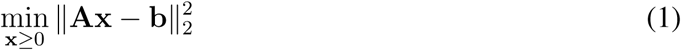

Each column of the matrix *A* represents a potential precursor to be considered as part of the observed spectrum. The observed peak intensities are represented by *b*. Each row of *A* then represents the library fragments that match the *m/z* of the corresponding row in *b*. If a precursor does not have a fragment that matches a particular peak, the entry in that row is a zero, while if it is a potential match, the entry is the normalized fragment intensity. To account for library fragments that do not match any peak in the spectrum, zero entries are added to the end of *b*; one for each fragment that has no corresponding match. This penalizes library spectra that have many fragments that are not matched to the spectrum. To encourage sparsity in the solution and account for the non-specificity of some ions between multiple channels of the same precursor, Elastic Net regularization is performed rather than plain NNLS. The L2 ratio is bounded between 0.1 and 0.9, and automatically determined per spectrum as the maximum cosine similarity between any two library precursors. The values of *x* that minimize Equation (1) are fed into an iteratively reweighted least squares (IRLS) problem, where over-predicted (less intense than expected) peaks that represent clean library intensity errors are used to compute a residual cutoff. This is then applied to the under-predicted (more intense than expected) peaks using an asymmetric Tukey’s bisquare loss that caps the influence of interfering fragment ions on the solution.

#### Scoring

During the main search, many attributes of the fitting process are collected in each scan for the respective precursors, both target and decoy. These include the retention time error, the fraction of the library intensity matched, and the scribe score^49^. A full list can be found in the Help folder of the GitHub repository. These attributes are only collected for candidate precursors in scans in which they were considered in the NNLS as described in the previous section. The data are then combined and condensed into a table of features, where each row corresponds to a candidate precursor identification. If performing plexDIA, these candidates are channel specific. Next, additional precursor properties, such as fragment correlations, are calculated for each candidate and added to the table. This table is then used to train the target-decoy model. By default, XGBoost^51^ is trained with 5-fold cross-validation to differentiate between targets and decoys. These folds are grouped such that predictions for each peptide are made with a model that has not been trained using that peptide in any charge state or channel. It also outputs the data used to train the models so that users can reanalyze their data, enabling the inclusion of additional features or external scoring models.

JMod uses the unbiased estimator for false discovery rate estimation proposed by Levitsky *et al.*^52^ for false discovery rate estimation. Q-values are estimated as the sum of the number of decoys+1 over the number of targets. When using non-isobaric mass tags, each channel is scored independently providing a *Qvalue* specific to that precursor in that channel. However, each precursor is also assigned a value *BestChannel Qvalue* which is the q-value of the best scoring channel in that plex-set. Protein q-values are calculated by assigning the maximum target or decoy score to each respective protein, before using the same methods.

#### Quantitation

Quantitation in JMod is achieved by performing additional rounds of NNLS fitting depending on the MS level. JMod provides significant flexibility in how quantification is performed at the MS1 level to make it suitable for a wide variety of labeling reagents. The apex of a precursor is first selected based on its maximum MS2 quantification (*coeff*), and in multiplexed experiments the channels will vote on a shared apex for quantitation. All channels and their theoretical isotopic envelopes are then fit to the observed signal in the spectra about this apex, giving a series of MS1 coefficients that define the MS1 elution peak of each channel. JMod returns the area under these peaks as *plex Area*.

The introduction of mass offsets with certain isotopes such as deuterium induces a retention time shift between the channels^35^. JMod enables different channel apex matching strategies to account for this. Channel apices can be forced into the same voted apex for reagents that exactly coelute, allowed to freely find their apex in the vicinity of the voted one, or allowed to walk monotonically to the most intense adjacent peaks.

By default, JMod reports the maximum coefficient predicted over all possible scans for MS2 quantitation (*coeff*). This is analogous to quantitation with a scan window range of 1. Users can also extract more intricate measures from the output files, as these include all predicted coefficients for each precursor in each spectrum.

#### Proteome Digest Sample Preparation

The two-proteome samples were prepared from human K562 proteins (Promega, V6941) and *S. cerevisiae* (yeast) peptides (Promega, V7461). Intact proteins were provided in a buffered urea solution to which dithiothreitol then iodoacetamide were added to reduce and alkylate cysteine residues per manufacturer’s recommendation. The urea was diluted with 120 mM ammonium bicarbonate (ABC) then digested with trypsin overnight at 37*^◦^*C using a 1:50 enzyme:protein ratio. Resulting peptides were desalted using C18 cartridges on a negative pressure manifold. Human and yeast peptides were added at quantitative ratios as demonstrated in Supplementary Table 2 for diethyl labeling. Alternatively, predigested human peptides (Promega, V6951) were combined with yeast peptides at ratios shown in Supplementary Tables 1 and 3 for PSMtag labeling.

#### Diethylation

The diethylation protocol was adapted from previous reports^53–55^. Dried peptides were reconstituted in 100 mM HEPES buffer to 500 ng/*µ*L. Light (C_2_H_4_O), medium (^13^C_2_H_4_O), or heavy (C_2_D_4_O) acetaldehyde was added from a 20% v/v stock to final concentrations of 2.5%. Sodium cyanoborohydride (NaBH_3_CN) and sodium cyanoborodeuteride (NaBD_3_CN) were prepared fresh daily and added to final concentrations of 250 mM reducing agent. Isotopes of acetaldehyde and the reducing agents were combined to obtain six possible masses. The diethylation reaction proceeded for 1 hr at 37*^◦^*C with shaking at 1000 rpm. Labeling was quenched for 10 min with ABC at a final concentration of 500 mM. The final peptide solution was acidified with formic acid, cleaned using house-packed C18 spin tips, and then dried. The dry peptides were reconstituted with mobile phase A (MPA, 0.1% formic acid in water), combined, and subjected to LC-MS/MS analysis in either a 3-plex or 6-plex combination.

#### PSMtag Labeling

PSMtag employs NHS-ester chemistry to label primary amines present on peptide N-termini and lysine residues. Samples were each labeled with one of nine non-isobaric isotopologues of PSM-tag: the lightest form, Δ00, and eight heavier forms increasing in mass by 2 Da each up to Δ16. All PSMtag analogues were synthesized and provided in powder form by CreaGen, Inc. Tags were solubilized in anhydrous DMSO to a concentration of 24 mg/mL. An organic base, N,N-diisopropylethylamine (DIPEA), was combined with the tag stocks to final concentrations of 4% (approx. 218 mM) DIPEA and 12 mg/mL PSMtag in DMSO. This labeling solution was added to all dried peptides mixtures (see Supplementary Tables 1 and 3) at a ratio of 1.5:1 tag to peptide mass. The reaction was allowed to proceed at room temperature for 1-2 hr prior to quenching with amine-functionalized paramagnetic beads (Catalog ID MR-AMN002, ReSyn Biosciences). Beads were washed with water then three times with anhydrous DMSO to mitigate possible tag hydrolysis in the presence of aqueous solvents. The beads were suspended in DMSO and added directly to the labeled peptides using a mass ratio of 1:1 beads to tag and incubated for 30 min. Labeled peptides were eluted from the beads using 500 mM triethylammonium bicarbonate (TEAB). Samples were either directly diluted to 30% acetonitrile, 0.05% DDM, and 2% formic acid prior to LC-MS/MS analysis or dried and resuspended in the same conditions.

### Hepatocyte Sample Preparation

#### Animal husbandry and metabolic labeling

All procedures were approved by the Institutional Animal Care and Use Committee at Massachusetts General Hospital. Young (4-month) female C57BL/6 mice were obtained from the National Institute on Aging (RRID:SCR 007317). To metabolically label proteomes with ^13^C_6_-lysine, animals were fed a Lys+0 control diet (Mouse Express L-Lysine [^13^C_6_, 99%] feed ad lib, Cambridge Isotope Laboratories) for a 10-day acclimation period, then switched to the Lys+6 labeled diet ad lib (approximately 4 g/day consumed) for either 3 or 5 days. Euthanasia was performed by CO_2_ asphyxiation followed by cervical dislocation, and tissues were collected after PBS perfusion. Fresh liver was dissociated into single-cell suspensions using the TIGR Tissue Grinder and Dissociator per the manufacturer’s protocol.

#### Single-cell sample preparation

The mouse liver single cells were prepared using the nPOP method for single-cell proteomics^56,57^. Cells between 24 and 43 *µ*m in diameter with elongation below 2.2 were sorted from single-cell suspensions using the CellenOne, onto fluorocarbon-coated glass slides into 9-nL droplets of 100% DMSO. These were incubated overnight with 100 ng/*µ*L Trypsin Platinum (Promega VA9000) and 5 mM HEPES at 72% relative humidity. The following day, the peptides were chemically labeled with 20 nL of mTRAQ reagents Δ0, Δ4, or Δ8, dissolved in 100% DMSO and then 20 nL of 100 mM TEAB. The samples were then incubated for one hour, pooled using the CellenOne in a 50/50 solution of acetonitrile and water and dispensed into a 384 well PCR plate.

#### Bulk digest preparation for spectral library

Some of the single-cell suspensions were also used to prepare a bulk digest for the two mice. Cells were pelleted and resuspended at a concentration of 1000 cells/*µ*L in MS grade water. Cells were lysed by freezing to -80*^◦^*C and then heating to 90*^◦^*C for 10 minutes. The resulting lysate was then digested overnight at 37*^◦^*C using trypsin to a concentration of 20 ng/*µ*L and 8.5 pH TEAB buffer to a concentration of 100 mM.

1 *µ*g of digested bulk peptide samples were dried down and resuspended in 1 *µ*L of 100 mM TEAB then labeled by adding 0.33 *µ*L of mTRAQ Δ0, Δ4, or Δ8 reagent at a concentration of 1/20th unit per *µ*L. Samples were incubated for 2 hours at room temperature, quenched with hydroxylamine at 0.2%, dried down and resuspended in 2% formic acid and 0.015% n-dodecyl-*β*-D-maltoside (DDM) for LC-MS/MS injection.

### LC-MS Acquisition

#### Bulk DE-proteome analysis

For demonstration of quantitative accuracy and precision of JMod for plexDIA data, diethylated peptide samples were injected as a 4-Da spaced 3-plex or a 2-Da spaced 6-plex. At least 100 ng per channel were injected on an IonOpticks Aurora Ultimate C18 column using a Vanquish Neo LC coupled to an Exploris 480 Orbitrap MS (Thermo Scientific). The gradient began at 5% mobile phase B (MPB, 80% acetonitrile with 0.1% formic acid) and immediately began separation starting at 10% MPB to 36% over one hour using a flow rate of 200 nL/min. MPB increased to 95% and remained for 8 min before column reequilibration back to 5% MPB. Orbitrap resolutions of 120,000 and 30,000 were used for MS1 and MS2 scans, respectively. A variable window scheme with 60 MS2 scans ranging from m/z 380 to 1370 all overlapping by 0.5 Th; the first 38 windows were 8.5 Th wide, the following 9 were 17.5 Th, and the final 13 were 41.5 Th wide. The overall cycle time for MS2 windows was ∼3.5 s with MS1 scans acquired every 1.5 s. A constant normalized collision energy of 30% was applies with maximum injection time set to ”Auto” to be synchronized with Orbitrap acquisition duration.

### PSMtag data acquisition

#### Benchmarking quantitative performance

PSMtag-labeled samples with human and yeast ratios as outlined in Supplementary Table 1 were separated with an IonOpticks Aurora XT C18 column using a Vanquish Neo LC configured for direct injection. The gradient began at 30% MPB and separation occurred at 0.1 min starting from 38% MPB to 62% for 24.5 min and to 75% MPB for another 5.4 min using a flow rate of 100 nL/min. Peptides were ionized with an electrospray ionization voltage of 1900 V and introduced to an Orbitrap Astral MS equipped with a FAIMS Pro Duo source, which implemented a constant -50 V compensation voltage and 3.6 L/min gas flow rate. MS1 spectra were collected every 600 ms with a maximum injection time of 150 ms and resolving power of 240,000. MS2 spectra were collected using the Astral analyzer with 20 variably sized windows shown in 4 with a maximum injection time of 60 ms (∼1200 ms MS2 cycle time) and normalized collision energy of 24%.

#### Large fold change data

Large fold changes, as described in Supplementary Table 1, were generated to evaluate the effect of ratio compression. Samples were injected at various input amounts spanning 200 pg/channel to 20 ng/channel of the human reference on the Vanquish Neo - Orbitrap Astral system. The LC was configured for trap-and-elute using an IonOpticks NanoShield trapping column. The gradient began at 30% MPB at a flow rate of 300 nL/min before decreasing to 150 nL/min after 0.1 min, at which time separation started at 32% MPB and continued to 52% MPB for 24.5 min, then increased to 62% MPB for 5.4 min. MS1 spectra were collected with a maximum injection time of 20 ms, and resolving power of 240,000. To accommodate the difference in ion population for different input amounts, different MS/MS methods were employed for DIA acquisition. For single-cell equivalents at 200 pg/channel of human reference, 40 variably sized DIA windows were applied as described in Supplementary Table 5 with maximum injection times of 20 ms per window (∼800 ms MS2 cycle time). For the 2 and 20 ng/channel injections, fixed windows were used across the m/z range from 500 - 1000 Th using either 5- or 2-Th windows and 8- or 3-ms max injection times (∼800 and 750 ms MS2 cycle times), respectively. All other MS2 parameters were kept consistent across inputs, including normalized collision energy (24%) and MS1 cycle time (600 ms).

#### Acquisition of mouse liver single cells

Single cells acquired from mouse liver and labeled with mTRAQ were separated with a Vanquish Neo LC configured for trap-and-elute coupled to an Orbitrap Astral MS system (Thermo Scientific). nPOP-prepared samples were reconstituted in 2 uL, all of which was loaded onto a NanoShield trap and separated using an Aurora Ultimate C18 analytical column (IonOpticks). The gradient started at 6% MPB and increased to 29% in 15 min then increased again to 45% in another 15 min before washing the column with 99% MPB and reequilibrating back to 6% MPB. Full scans were acquired in the Orbitrap with a resolution of 240,000 and m/z range from 400-800. The Astral analyzer was used for all MS2 spectra which were collected using NCE of 28 with fixed 5-Th windows across the full m/z range using a maximum injection time of 10 ms (∼800 ms MS2 cycle time) and fixed MS1 cycle time of 600 ms.

### Data Analysis

#### Library generation

To provide a fair comparison, the same spectral libraries were used for both JMod and DIA-NN for each respective experiment. Similarly, the same library was used for JMod in both 2 Da and 4 Da spaced plexDIA comparisons.

Two sets of PSMtag-labeled data are included in this manuscript. The first dataset, presented in Fig. 2 and Fig. 3, contains mixed human-yeast samples collected in a 1-plex configuration with the Δ0 (light) channel only, as well as 5-plex and 9-plex variants. The second dataset, presented in Fig. 4, contains only 5-plex and 9-plex variants collected at three different input amounts. Due to the unique characteristics of the tag, preliminary human-yeast PSMtagged spectral libraries for the entire human and yeast proteomes were created using a publicly available fine-tuned retention time and fragmentation prediction model in DIA-NN 2.5.0. The PSMtag model was pretrained using DIA-NN 2.5.0^58^. Peptides from length 5 to 30 and charge states 1 to 5 were considered. To generate a library for the first set of data, the three Δ0 replicates were searched against the human-yeast PSMtag predicted libraries to generate a Δ0 empirical library. The resulting empirical library was then used to search the same data in JMod as well as the 5-plex and 9-plex data from the same dataset. This behaves similar to the second, match-between-runs search in DIA-NN only when searched against the same data that it was generated for, that is, the Δ0 data. For the second dataset, the largest input (20ng) 5-plex data were searched against DIA-NN to produce an empirical library. This search used the following flags to define modifications: {*–channels tag,d0,nK,0:0; tag,d4,nK,4.01096:4.01096; tag,d8,nK,8.026839:8.026839; tag,d12,nK,12.033938:12.033938; tag,d16,nK,16.038574:16.038574 –original-mods*}. That was then used for the three 9-plex input amounts (20ng, 2ng, 200pg) and smaller input amounts for the 5-plex sample (2ng and 200pg). Again, when searched against the same 20ng 5-plex samples, it will behave like an MBR search rather than an empirical library.

The human-yeast diethyl labeled library was created using DIA-NN 2.5.1. In silico trypsin digests of the human and yeast fasta proteomes were used. Peptides with a maximum missed cleavage of one, length 7 to 30, and charge states 1 to 4 were predicted. The following flags were used to define modifications: {*–fixed-mod Diethyl,56.06260026,nK –lib-fixed-mod Diethyl – channels Diethyl,0,nK,0:0; Diethyl,4,nK,4.01342:4.01342; Diethyl,8,nK,8.05021:8.05021*}. This was then searched against the 3-plex diethyl data. The resulting empirical library from this search was used for all subsequent diethylation analyses. The label-free library was generated in the same way from bulk label-free data with similar parameters and without the inclusion of the diethly modification.

mTRAQ libraries for the mouse analysis were generated from labeled bulk injections of the murine liver samples. These data were searched with DIA-NN using a DIA-NN mTRAQ predicted library of the *M. musculus* proteome fasta. The following flags were used to define modifications for mTRAQ with heavy and light lysine: {*–fixed-mod mTRAQ,140.0949630177,nK –channels mTRAQ,0L,nK,0:0; mTRAQ,0H,nK,0:6.0201290268; mTRAQ,8L,nK,8.0141988132:8.0141988132; mTRAQ,8H,nK,8.0141988132:14.03432784*}. The resulting spectral library was used for all mouse liver analysis by JMod.

#### Entrapment

The entrapment FDRs were calculated using the paired method from Wen et al^31^. Briefly, preliminary libraries were predicted from full fastas as described in Library generation. As both multiplexing and entrapment multiplicatively increase the library size, these predicted libraries were filtered to approximately 150K precursors, much larger than the number that would be identified, to allow reasonable search times. For each target sequence in an input library, the peptides are grouped by modified sequences, then shuffled by permuting all but the last amino acid in the target sequence with the modified sequence as a seed. If after 50 shuffling attempts no entrapment sequence was found that was unique from the original target sequences and was not already present in the generated entrapment sequences, both that target and the corresponding entrapment sequence were removed from the database. There was therefore a one-to-one correspondence between the original targets and their shuffled entrapment sequences in the resulting library. Retention times were predicted for both target and entrapment sequences using JMod’s internal retention time prediction models to maintain parity between the two sets of sequences. The entrapment sequence fragment *m/z*s were adjusted according to the permutation at the same fragment indices, while the intensities remained the same. For the paired entrapment method, Wen et al. proposed the following formula:^31^

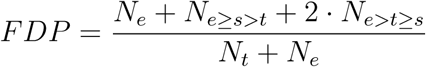

where *N_e_* and *N_t_*, respectively, represent the number of entrapment targets and original targets scoring better than the current score threshold, *s*. *N_e≥s>t_*indicates the number of entrapment targets at *s* for which their scores are greater than or equal to *s* but the score of their respective original complements are lower than *s*. Likewise *N_e>t≥s_* indicates the number of entrapment targets for which their scores were greater than their respective original targets, and for which their original targets also scored greater than or equal to *s*.

#### Search parameters

For all label-free analysis, JMod 2.0.0 was used with *–no ms1 req* and otherwise default parameters.

In the label-free entrapment analysis only, DIA-NN 1.9 was used, as this version provides internal scores (CScores) that can be used to calculate the entrapment FDR. For these analyses the data were searched with match between runs turned off, run-to-run normalization disabled, protein inference turned off, the precursor FDR(%) set to 50, and with otherwise default parameters. Match between runs was turned off so that replicates were searched independently to allow fair evaluation between the software. In other label-free analysis, DIA-NN 2.5.1 was used with default parameters and quantitation set to *–high-acc*.

All PSMtag data was searched using JMod 2.0.0 with *–plexDIA* and either *–tag PSMtag*, *– tag PSMtag 5plex* or *–tag PSMtag 9plex* labels applied, respectively. To account for slight drift between channel apices, we also allowed quantitation to be performed within ± 3 scans of the shared apex with with the *–additional scans 3 –free apex* flags. All other parameters were left at the JMod defaults with the exception of *–no ms1 req*.

DIA-NN version 2.5.1 was used to search the PSMtag data. PSMtag was added as a fixed modification using {*–fixed-mod tag, 308.1160923903, nK –channels tag,d0,nK,0:0; tag,d4,nK,4.01096:4.01096; tag,d8,nK,8.026839:8.026839; tag,d12,nK,12.033938:12.033938; tag,d16,nK,16.038574:16.038574 –original-mods –lib-fixed-mod tag*}. Quantitation was set to *–high-acc*. The q-value threshold was set at 0.01. Protein inference was disabled. The normalization strategy was set to channel-spec-norm.

JMod version 2.0.0 was used for all diethylation searches. The data were searched using either *–tag diethyl 3plex* or *–tag diethyl 6plex* respectively. Due to the larger retention time shifts associated with diethylation, the parameters *–additional scans 20 –free apex* were also used. Again, other parameters were left at the JMod defaults with the exception of *–no ms1 req*.

For the diethylation searches, DIA-NN version 2.5.1 was used. The same parameters as the PSMtag searches were used with channels defined with the following commands; {*–fixed-mod Diethyl, 56.06260026, nK –lib-fixed-mod Diethyl –channels Diethyl,0,nK,0:0; Diethyl,4,nK,4.01342:4.01342; Diethyl,8,nK,8.05021:8.05021; –original-mods*}

#### Quantitative comparisons

For quantitative comparisons, MS1 quantities for DIA-NN are shown using the *Ms1.Normalized* of precursors. QuantUMS values are taken from the *Precursor.Normalized* column. We restrict the precursors reported by DIA-NN in this analysis to those with both a *Q.Value<*0.01 and *Channel.Q.Value<*0.01 so that DIA-NN has its highest quality identifications included for comparison. For JMod, MS1 quantitation is shown using *plex Area* for precursors with *Qvalue<*0.01. All comparisons for precursor-level quantification are show on the intersected set of precursors detected by DIA-NN in the 5-plex data with values for both MS1 and QuantUMS quantification, JMod in the 5-plex data, and JMod in the 9-plex data. All cross-channel comparisons are done after recentering on the median human fold changes between the channels. For the comparisons done in Fig. 4, all cross-channel comparisons for a given expected log_2_FC were computed in Fig. 4, whereas in Fig. 3 all channels were compared only to the reference (maximum yeast input amount) channels to aid in interpretability due to small fold-change spacing between the input amounts.

For comparisons in the number of IDs (precursors, proteins), different filters were used. JMod only produces run-level precursor and protein q-values, whereas DIA-NN produces run- and channel-level precursor q-values but only channel-level protein q-values. As such, it is difficult to make direct comparisons in the number of IDs. For precursors counts we use DIA-NN’s *Q.Value* column, where q-values are essentially identical across different channels in the plex, and filter to *<*0.01, and for JMod we filter based on the *BestChannel Qvalue* column. DIA-NN does not report run-level protein q-values in plexDIA mode, so we attempt to approximate it by computing the union of protein IDs passing a q-value filter of *<*0.01 in each channel from the *PG.Q.Value* column. For JMod, we filter the *Protein Qvalue* column to *<*0.01. Note that this metric may be artificially conservative for DIA-NN compared to JMod and that if it were to report an equivalent metric then reported protein IDs may be higher.

#### Hepatocyte Analysis

Hepatocytes were analyzed using the QuantQC package^41^ similar to Leduc *et al.*^22^. In brief, failed cells first were excluded using total protein estimates from negative controls. A peptide-cell matrix was then created using at most 5 peptides per protein. The matrix was then normalized to relative *log*_2_ fold changes^59^. The effects of LC runs were subtracted from each peptide where the run effect spline had an *R*^2^ *>* 0.1. The data were then collapsed into a protein-cell matrix by taking the median intensity of peptides mapping to each protein. KNN imputation was used before batch correction for label and mouse. Finally, erythrocyte contamination was removed by subtracting out signal correlating with P01942 (Hemoglobin subunit alpha).

Protein degradation rates were also calculated according to Leduc *et al.*^22^. In summary, initial estimates for degradation rates were calculated using *log*(*Heavy/Light* + 1)*/t*. The fraction of heavy lysine in the free amino acid pool for both mice was then estimated from bulk samples. This was then used to correct the degradation rates of the proteins as amino acids can come from the feed or from amino acids mobilized from degraded proteins^60^.

To confirm the axis of zonation, protein abundances were downloaded from Fig. 5**a** of Ben-Moshe *et al.*^36^ Portal proteins were selected as the top 10 with their center of mass closest to the portal vein according to Ben-Moshe *et al.* with p-values less than 0.01 in both our data and theirs. Central proteins were selected in a similar way with the top ten closest to the central vein. PC1 was divided into 9 equal partitions to help plot the data. Median protein abundances for cells in each partition were then plotted for the 20 zonation markers RNA abundances were downloaded from Figure 3**c** of Halpern *et al.*^38^. The proteins for Extended Data Fig. 6c were selected as follows. Proteins were filtered for those present in at least 300 cells. Degradation regulated proteins were selected as the 10 where degradation and abundance were most negatively correlated with one another and negatively correlate across PC1. Similarly, RNA regulated proteins were selected as the 10 where RNA abundance and protein abundance most positively correlate with one another.

Data for Extended Data Fig. 6e was filtered as follows. Cells were divided into 3 groups (portal, neutral, and central), using PC1, where higher values indicate central cells (Extended Data Fig. 6a). Protein half lives were then calculated as *log*(2)*/K_d_* where *K_d_* is the degradation rate of the protein. Proteins were then filtered for those with measurable half lives in at least 50% of portal cells and 50% of central cells. All code to recreate the analysis can be found at github.com/ParallelSquared/JModFigures.

## Extended Data Figures

**Extended Data Fig. 1.**
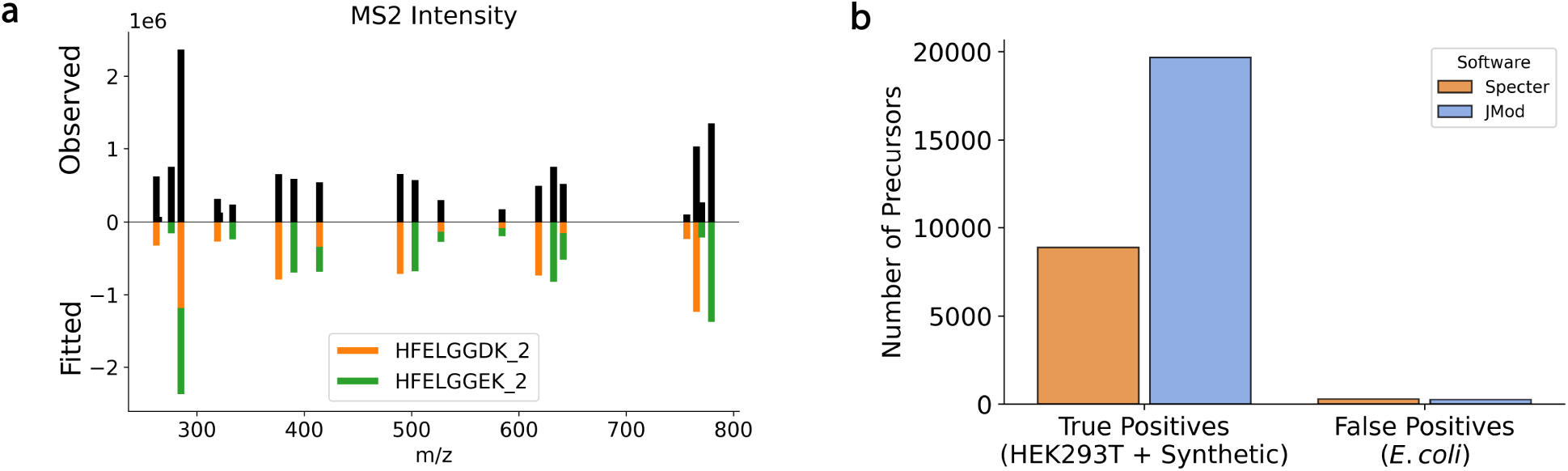
Overlapping peaks in MS2 spectra. **a** Mirror plot of the observed and fitted MS2 fragment intensities for 2 coeluting precursors; (HFELGGDK, 2+) and (HFELGGEK, 2+). While JMod also models the isotopic distributions of each fragment, only peaks that match the monoisotopic fragments of the precursors are shown to enhance clarity. **b** Comparison of JMod with Specter^25^, a previous DIA software that modeled overlapping peaks in MS2 spectra. JMod was evaluated on the data from the Specter manuscript and compared to the Specter results reported therein.

**Extended Data Fig. 2.**
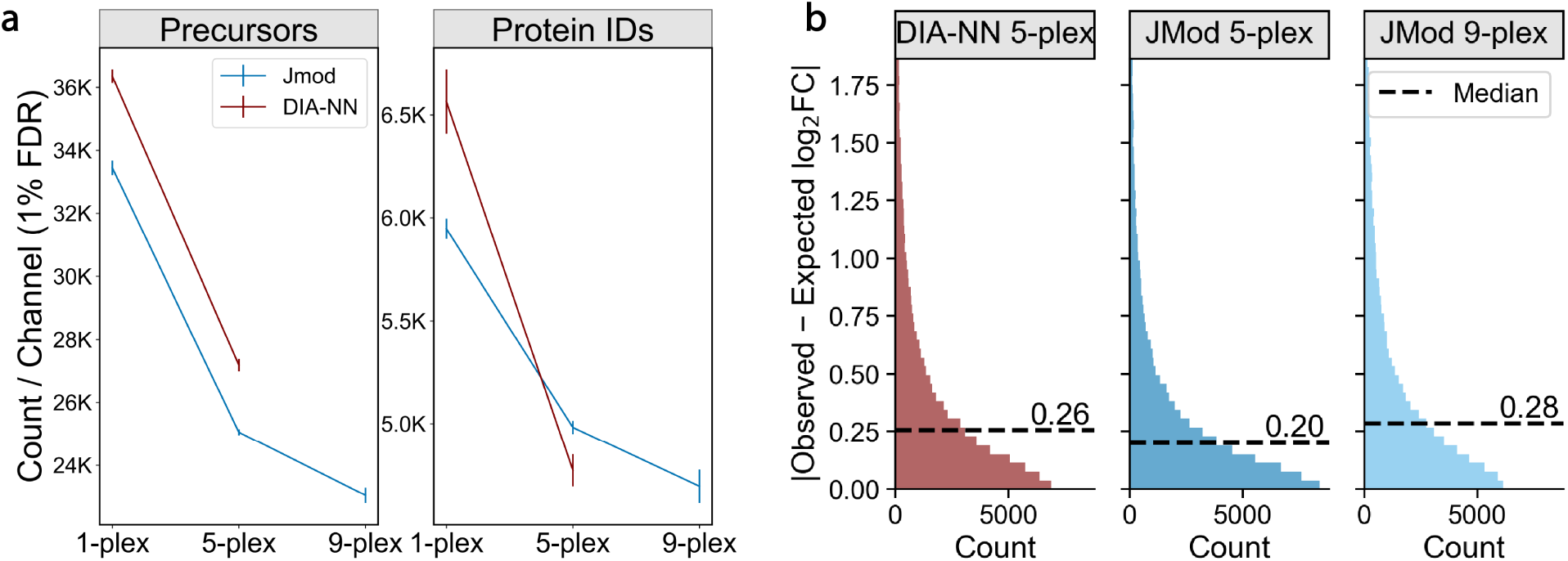
Comparing ID rates and quantitative accuracy. **a** Line plots with error bars showing standard deviations around the number of human precursor and protein identifications per channel for JMod and DIA-NN on single cell level PSMtag samples. **b** Histograms of mean absolute errors between the expected and estimated fold changes for intersected human precursors between DIA-NN 2.5.1 and JMod for 5-plex PSMtag data and 9-plex JMod PSMtag data.

**Extended Data Fig. 3.**
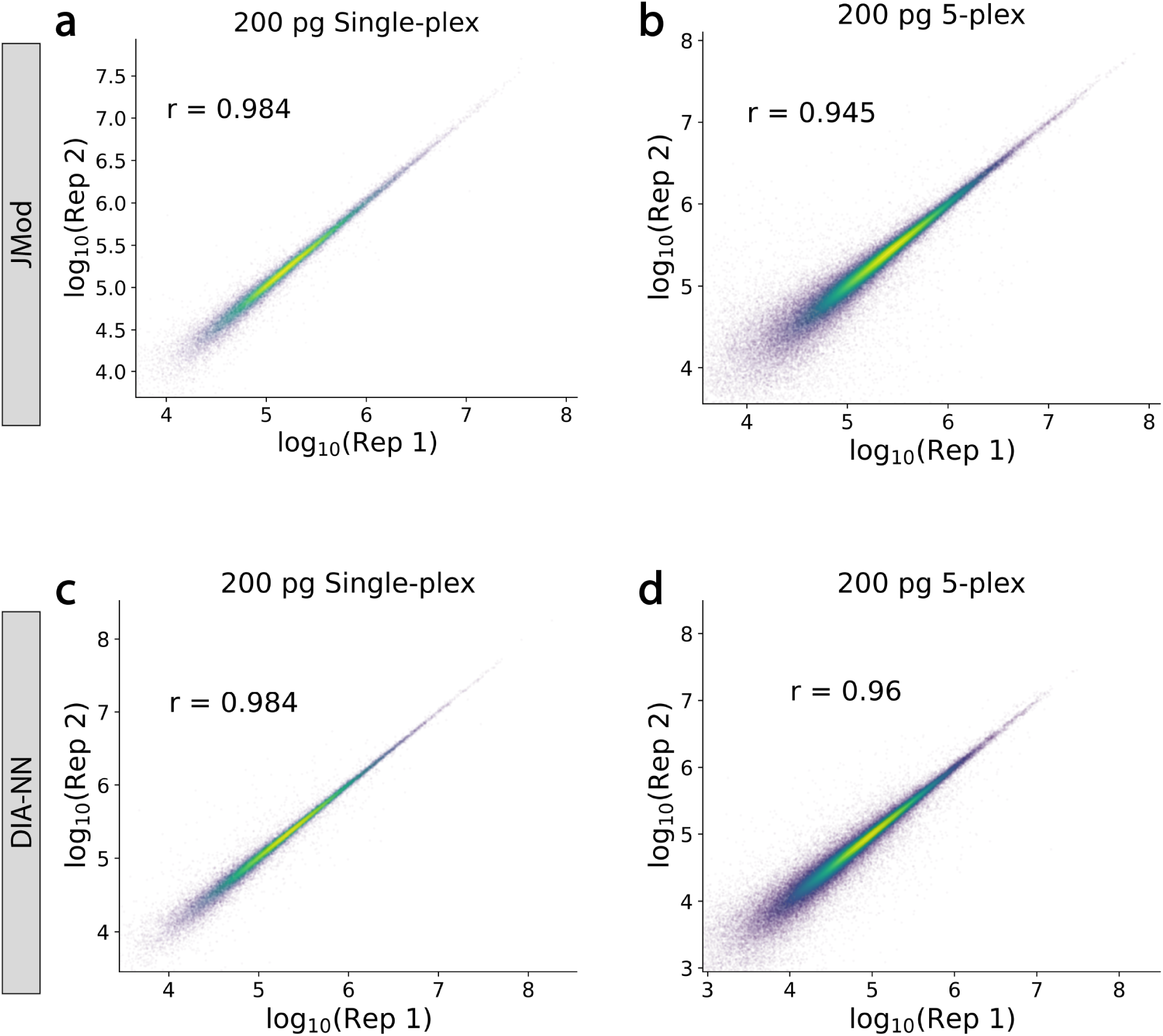
Comparing precursor quantities from 200 pg per channel replicates. **a** Comparison of two single-plex JMod replicates. **b** Comparison of two 5-plex JMod replicates. **c** Comparison of two single-plex DIA-NN 2.5.1 replicates. **d** Comparison of two 5-plex DIA-NN 2.5.1 replicates.

**Extended Data Fig. 4.**
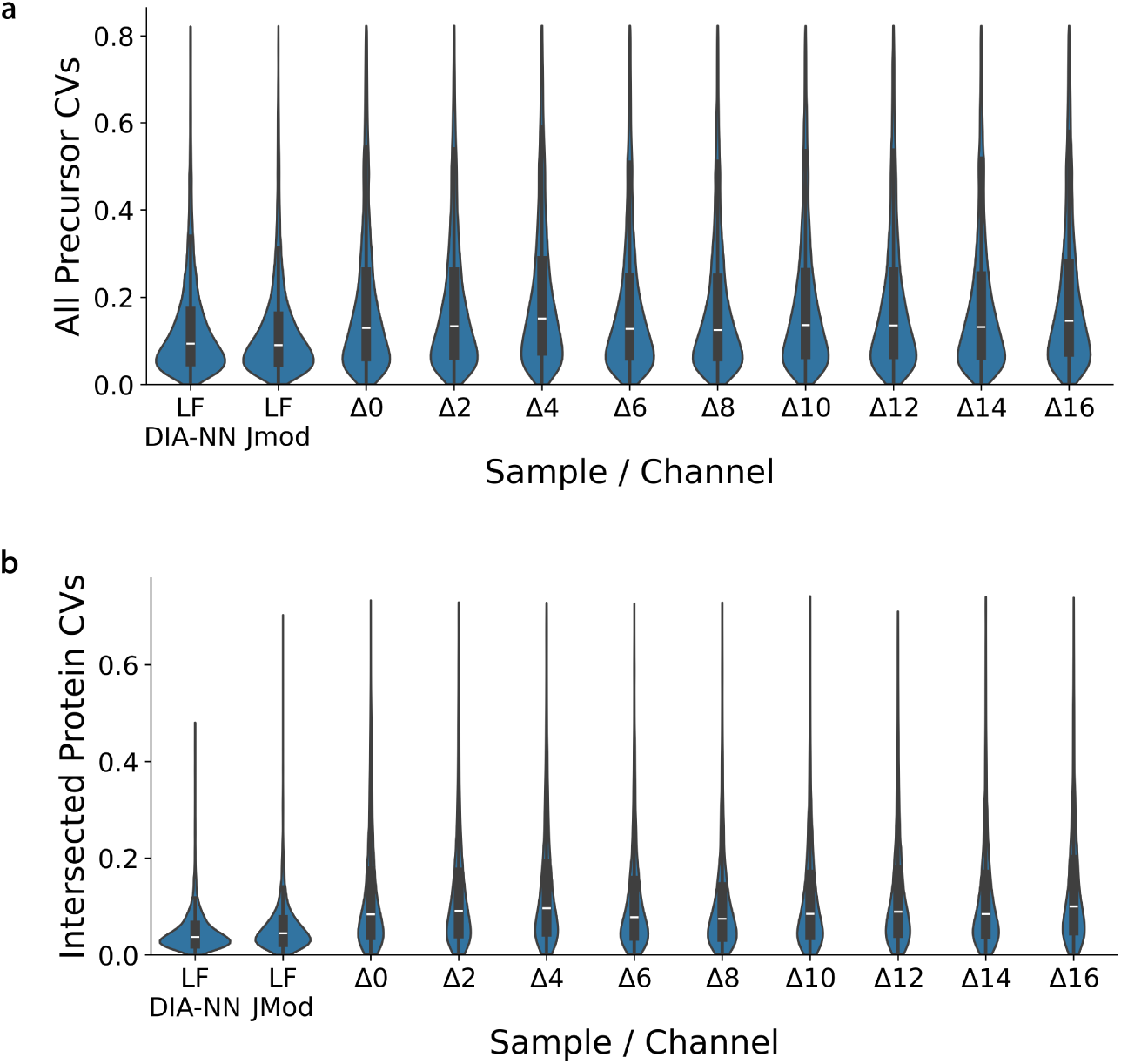
Evaluating CVs. **a** Comparison of precursor coefficients of variation (CVs) for all IDs from DIA-NN 2.5.1 and JMod on label free data as well as JMod on 9-plex PSMtag data. CVs were calculated from three replicates. Only data below the 95th percentile are shown. **b** Comparison of protein CVs for intersected IDs from DIA-NN 2.5.1 and JMod on the same PSMtag data. Only proteins with at least 2 confidently identified precursors were considered. Only data below the 99th percentile are shown.

**Extended Data Fig. 5.**
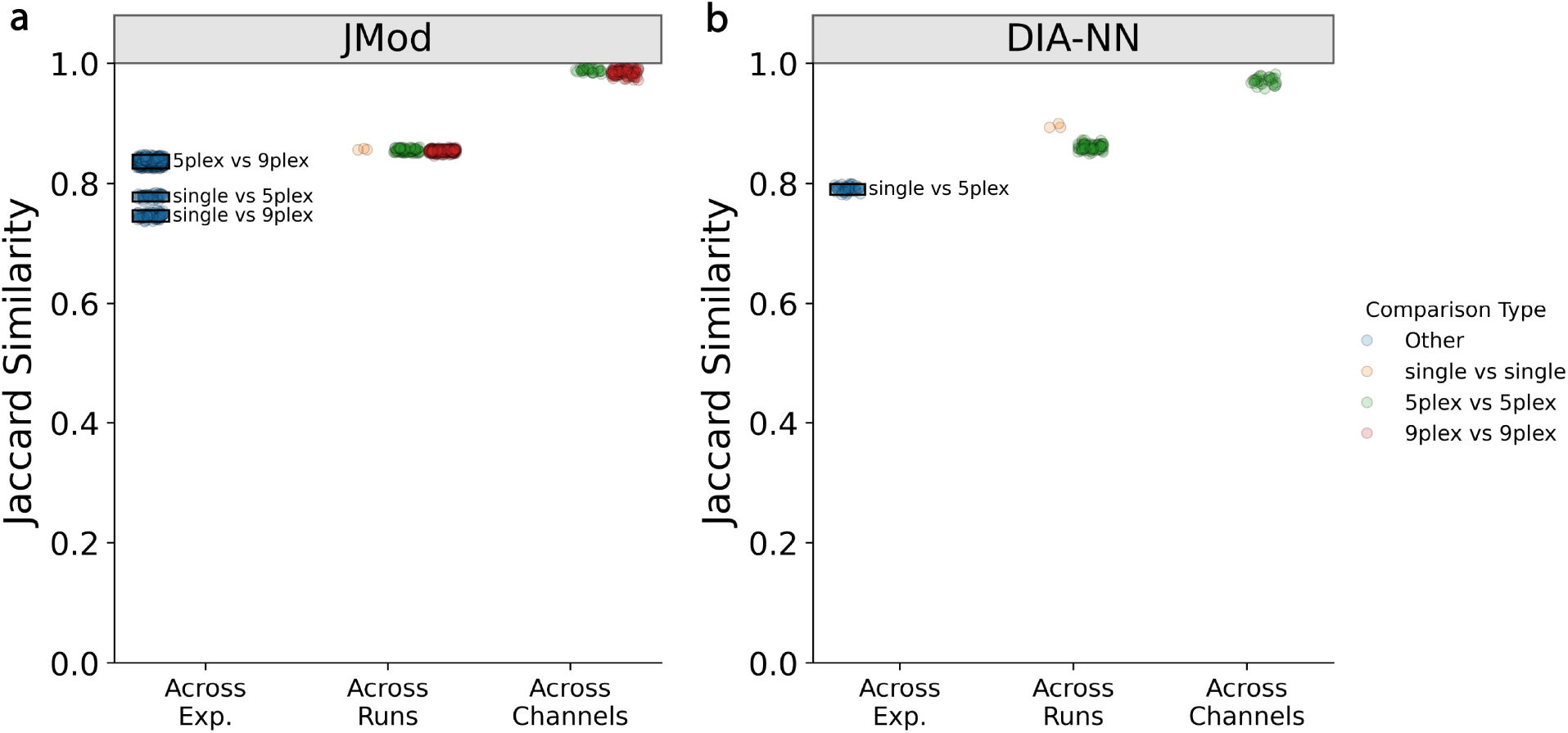
Consistency of plexDIA identifications. **a** Comparison of JMod protein identifications across channels (same plex-size), across runs (same plex-size), and across experiments (different plex-sizes). Data were PSMtag labeled samples at input amounts of 200 pg per channel. **b** Similar comparison for DIA-NN 2.5.1 protein identifications on the same data.

**Extended Data Fig. 6.**
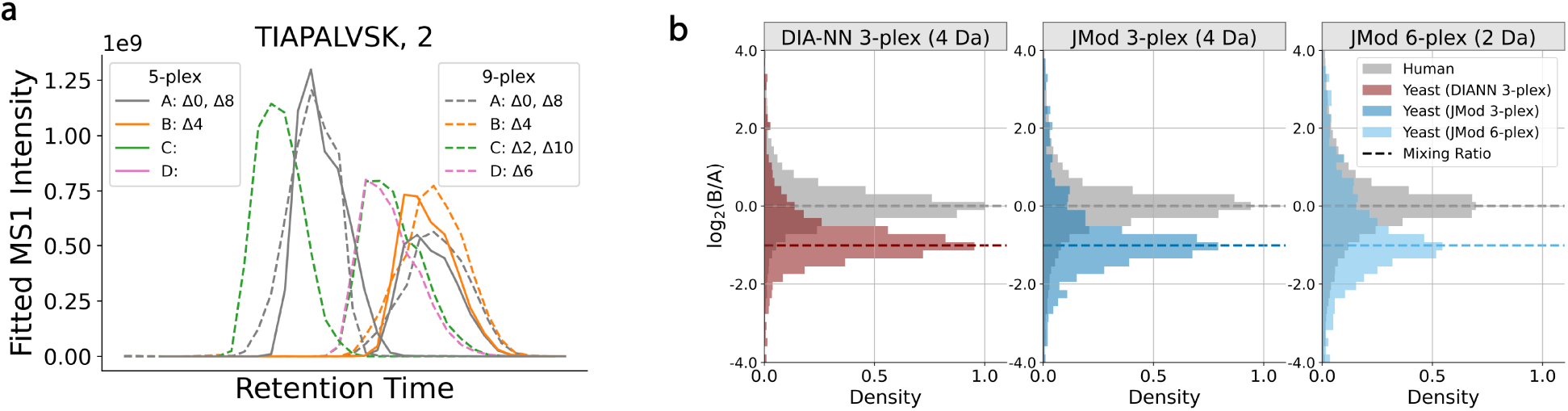
Quantitation of diethylated samples. **a** JMod’s fitted MS1 coefficients for 3-plex and 6-plex diethylated samples for Human peptide TIAPALVSK. **b** Distributions of fold change estimates for intersected human and yeast precursors from DIA-NN 2.5.1 and JMod for 3-plex diethylated samples and JMod only for 2 Da spaced 6-plex diethylated samples. Data are from the Δ0 and Δ4 channels with no retention time separation. Expected ratios are shown by colored dashed lines. Quantification by DIA-NN used the column *Precursor.Normalized* while quantification by JMod used the column *plex Area*.

**Extended Data Fig. 7.**
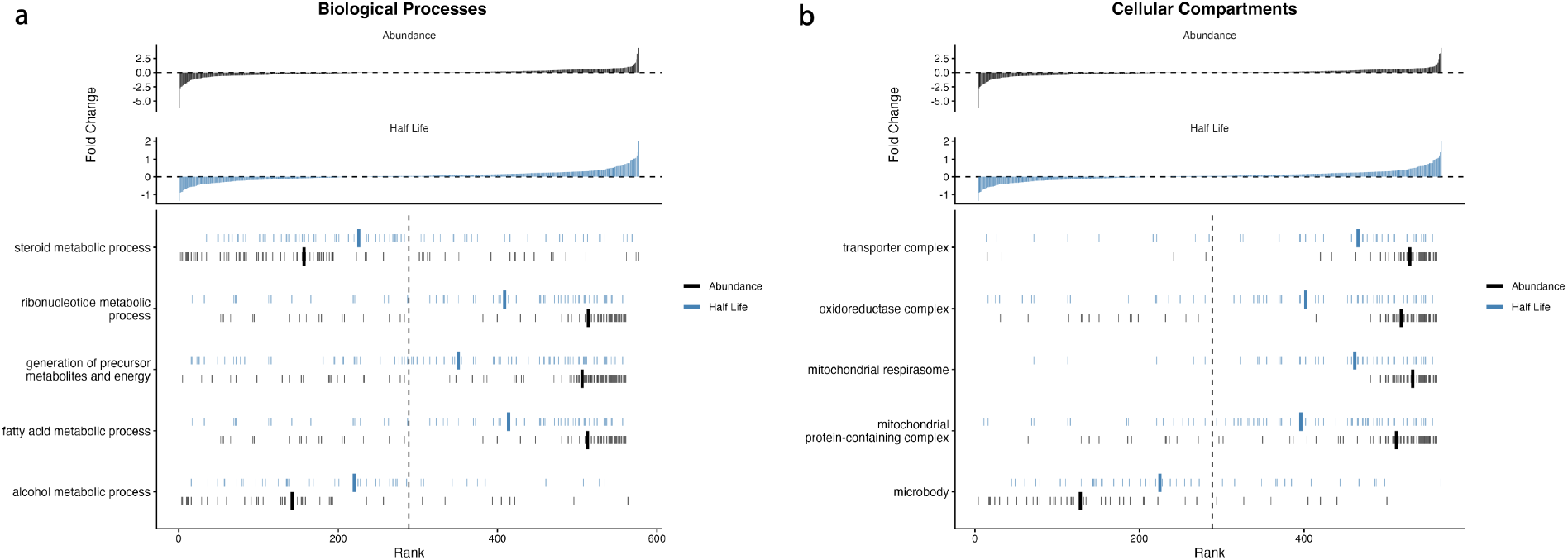
Significantly Zonated GO-Terms. **a** Biological processes where both protein abundance and degradation were significantly zonated. **b** Cellular compartments where both protein abundance and degradation were significantly zonated.

**Extended Data Fig. 8.**
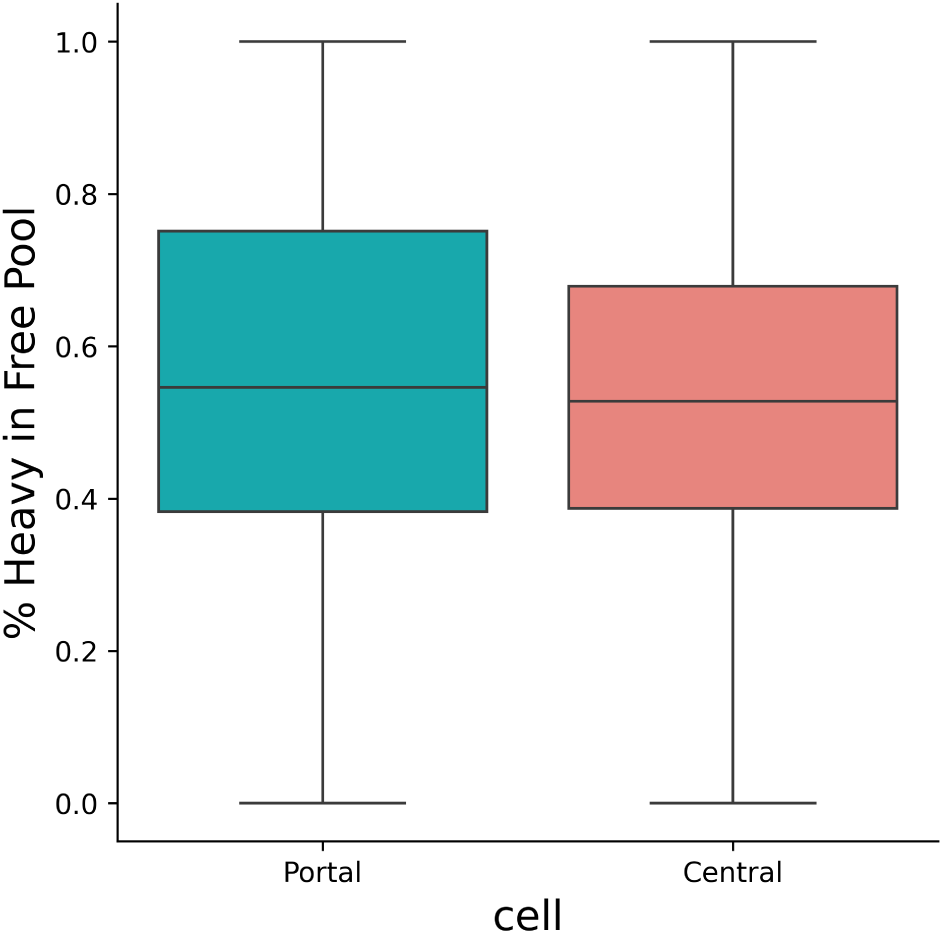
% Heavy in the Free Pool. Comparison of the percent heavy lysine available in the free amino acid pools for portal and central cells.

## Supplementary Information

**Supplementary Table 1.**
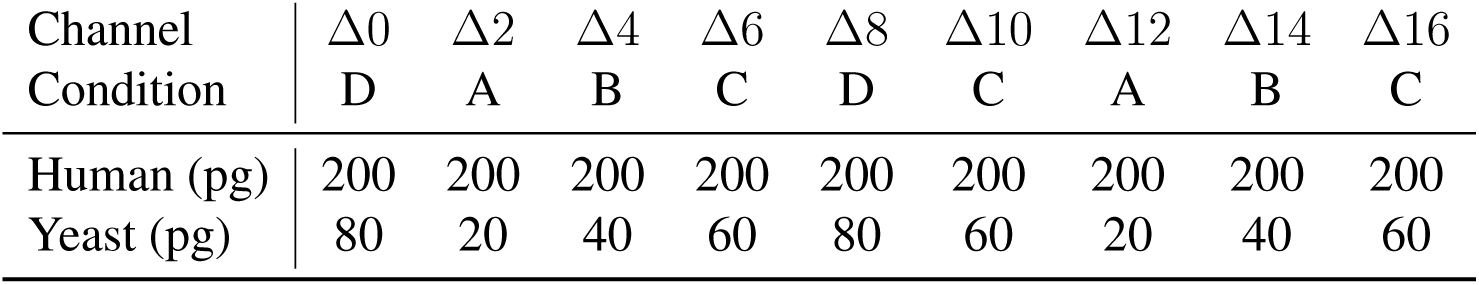
Mixing ratios for PSMtag two-proteome experiment.

| Channel<br>Condition | $\Delta 0$<br>D | $\Delta 2$<br>A | $\Delta 4$<br>B | $\Delta 6$<br>C | $\Delta 8$<br>D | $\Delta 10$<br>C | $\Delta 12$<br>A | $\Delta 14$<br>B | $\Delta 16$<br>C |
| --- | --- | --- | --- | --- | --- | --- | --- | --- | --- |
| Human (pg) | 200 | 200 | 200 | 200 | 200 | 200 | 200 | 200 | 200 |
| Yeast (pg) | 80 | 20 | 40 | 60 | 80 | 60 | 20 | 40 | 60 |

**Supplementary Table 2.** Mixing ratios for diethyl labeled two-proteome experiment.

| Channel<br>Condition | $\Delta 0$<br>A | $\Delta 2$<br>C | $\Delta 4$<br>B | $\Delta 6$<br>D | $\Delta 8$<br>A | $\Delta 10$<br>C |
| --- | --- | --- | --- | --- | --- | --- |
| Human (ng) | 100 | 100 | 100 | 100 | 100 | 100 |
| Yeast (ng) | 6 | 18 | 12 | 24 | 6 | 18 |

**Supplementary Table 3.** Mixing ratios for high fold-change PSMtag two-proteome experiment.

| Channel<br>Condition | $\Delta 0$<br>E | $\Delta 2$<br>A | $\Delta 4$<br>D | $\Delta 6$<br>B | $\Delta 8$<br>E | $\Delta 10$<br>C | $\Delta 12$<br>A | $\Delta 14$<br>C | $\Delta 16$<br>B |
| --- | --- | --- | --- | --- | --- | --- | --- | --- | --- |
| Human (pg) | 200 | 200 | 200 | 200 | 200 | 200 | 200 | 200 | 200 |
| Yeast (pg) | 640 | 20 | 320 | 80 | 640 | 160 | 20 | 160 | 80 |

**Supplementary Table 4.** 20 Window DIA Method.

| Label-free | PSMtagged |
| --- | --- |
| 400-438.5 | 500-538.5 |
| 438-466.5 | 538-566.5 |
| 466-490.5 | 566-590.5 |
| 490-511.5 | 590-611.5 |
| 511-533.5 | 611-633.5 |
| 533-549.5 | 633-649.5 |
| 549-563.5 | 649-663.5 |
| 563-574.5 | 663-674.5 |
| 574-585.5 | 674-685.5 |
| 585-600.5 | 685-700.5 |
| 600-616.5 | 700-716.5 |
| 616-628.5 | 716-728.5 |
| 628-643.5 | 728-743.5 |
| 643-654.5 | 743-754.5 |
| 654-671.5 | 754-771.5 |
| 671-689.5 | 771-789.5 |
| 689-716.5 | 789-816.5 |
| 716-747.5 | 816-847.5 |
| 747-793.5 | 847-893.5 |
| 793-900 | 893-1000 |

**Supplementary Table 5.** 40 Window DIA Method.

| PSMtagged |
| --- |
| 500-517.5 |
| 517-538.5 |
| 538-554.5 |
| 554-566.5 |
| 566-578.5 |
| 578-590.5 |
| 590-600.5 |
| 600-611.5 |
| 611-622.5 |
| 622-633.5 |
| 633-643.5 |
| 643-649.5 |
| 649-654.5 |
| 654-663.5 |
| 663-667.5 |
| 667-674.5 |
| 674-679.5 |
| 679-685.5 |
| 685-693.5 |
| 693-700.5 |
| 700-709.5 |
| 709-716.5 |
| 716-722.5 |
| 722-728.5 |
| 728-736.5 |
| 736-743.5 |
| 743-749.5 |
| 749-754.5 |
| 754-761.5 |
| 761-771.5 |
| 771-780.5 |
| 780-789.5 |
| 789-801.5 |
| 801-816.5 |
| 816-833.5 |
| 833-847.5 |
| 847-867.5 |
| 867-893 |
| 893-929 |
| 929-1000 |

